# Lifestyles of *Gypsy*-family transposons shape their regulatory mechanisms

**DOI:** 10.64898/2026.05.19.726053

**Authors:** Anna-Maria Papameletiou, Benjamin Czech Nicholson, Susanne Bornelöv, Gregory J. Hannon

## Abstract

Transposable elements are a highly diverse group of selfish genomic elements, prevalent across the tree of life, whose uncontrolled propagation poses a threat to genome stability. Recent studies have explored the evolution of *Drosophila melanogaster* transposable elements, their co-evolution with the host genome, and mechanisms that regulate their activity. However, little is known about their cross-species evolutionary patterns. Long terminal repeat (LTR) retrotransposons are the most active group of transposable elements in *Drosophila*. They are broadly separated into retroelements, which are active in the germline, and insect endogenous retroviruses that express in the gonadal soma. Somatic elements are hypothesised to infect the germline through their acquisition of virus-derived proteins such as Envelope and sORF2, thus multiplying through successive generations. In this study, we curated the sequences of LTR retrotransposons in 249 drosophilid genomes, allowing us to study their evolution across these species and highlight their varying degrees of conservation. Furthermore, we reveal multiple instances of Envelope protein loss or inactivation that suggest shifts in the expression pattern of these transposons, likely accompanied by adopting different transcriptional control mechanisms. We contrast this with the evolutionary history of sORF2, which we found to be much more stable. Lastly, we examined variations in transposon LTR regions responsible for transcriptional regulation and use predictive modelling to suggest six transcription factors likely involved in their tissue-specific expression. Altogether, we reveal complex, interspecies evolutionary patterns of *Gypsy*-family LTR retrotransposons and highlight examples of their coevolution with their host genome.

## INTRODUCTION

Transposable elements (TEs) are mobile genetic elements that threaten to compromise genomic integrity. TEs constitute roughly 20% of the *Drosophila melanogaster* genome, 45% of the human genome, with much higher burden observed in other eukaryotes such as salamanders and some plants (Xu *et al*, 2024; Decena-Segarra & Rovito, 2024). The interplay between TEs and host genomes is complex. TEs propagate by eluding host silencing mechanisms and, consequentially, the host adapts to counteract those challenges. This can be described as an evolutionary arms race and this ongoing molecular battle drives evolution, resulting in the rapid diversification of TE-related sequences both within and across species.

TEs are categorised according to their transposition mechanism (Mérel *et al*, 2020). Of particular interest are *Gypsy*-family long terminal repeat (LTR) retrotransposons that move via a “copy-and-paste” mechanism and are the most prominent elements in the *D. melanogaster* genome (Barckmann *et al*, 2018). These TEs are further subdivided according to their evolutionary relationships and include *Gypsy/gypsy (Metaviridae), Gypsy/mdg1, Gypsy/mdg3* and *Gypsy/chouto* (Kapitonov & Jurka, 2003; Nefedova & Kim, 2009).

All *Gypsy-*family LTR retrotransposons contain two identical LTRs surrounding an internal region containing open reading frames (ORFs) for Group-specific Antigen (*GAG*), which encodes structural proteins, and Polymerase (*POL*), which contains activities facilitating reverse transcription and integration (Hunter, 1994). An interesting characteristic of the *Metaviridae Gypsy/gypsy* clade is the acquisition of cell-to-cell movement capabilities, which splits the clade into retroelements and insect endogenous retroviruses. The latter contain an additional ORF encoding Envelope (ENV), which enables soma to germline infectivity by *gypsy* (Kim *et al*, 1994; Pelisson *et al*, 1994; Song *et al*, 1994; Keegan *et al*, 2021) and other TEs from the *Gypsy/gypsy* clade of which *gypsy* is a member (Marsano *et al*, 2000; Leblanc *et al*, 2000). The Envelope protein shares evolutionary origin with *Baculoviruses* (Malik et al., 2000; Rohrmann & Karplus, 2001; Terzian et al., 2001) and is thought to only have been acquired once by an ancestral *Drosophila* transposon (Senti *et al*, 2025; van Lopik *et al*, 2023).

In endogenous retroviruses (ERV), the F-type glycoprotein allows the formation of virus-like particles (VLPs), allowing the spread of ERV content amongst different cell types via endocytosis (Checkley *et al*, 2011; Brasset *et al*, 2006; Senti *et al*, 2025). More recently, sORF2, a fusion associated small transmembrane (FAST)-like protein, has been implicated in soma-to-germline transfer of all four *Gypsy/mdg1* family transposons (Voichek *et al*, 2025). FAST proteins facilitate viral spread by mediating membrane fusion, enabling cell-to-cell transfer without an extracellular phase (Ciechonska & Duncan, 2014). sORF2, in particular, has been shown to form channels between cells to facilitate transfer of TE material (Voichek *et al*, 2025).

TEs, like other genomic content, can be vertically transmitted from one generation to the next. However, their evolutionary histories are often much more complex due to introgression (Haring *et al*, 1995) which is widespread in the *Drosophila* evolutionary history (Suvorov *et al*, 2022) and horizontal transposon transfer (HTT). HTT is common in insects, including members of the *Drosophila* clade, where up to 55% of the total genome may have arisen as an indirect result of HTT (Pianezza & Kofler, 2025). HTT has been proposed to occur via direct transfer through a VLP (Brasset *et al*, 2006), piggybacking on exogenous vectors such as viruses (Miller & Miller, 1982) and parasitic mites and wasps (Houck et al, 1991; Yoshiyama et al, 2001). Additionally, parasitic bacteria called *Wolbachia* have been observed to horizontally transfer genes amongst *Drosophila* species (Dunning Hotopp et al, 2007), and similar transfer of TEs has been hypothesised (Loreto *et al*, 2008). Both introgression and HTT enables TEs to invade new species: in total, out of 122 well-characterised *Drosophila melanogaster* TEs, an estimated 11 have invaded the species in the past 200 years (Pianezza & Kofler, 2025). The majority of these come from species within the *Sophophora* subgenus, *D. simulans* and *D. willistoni* (Pianezza *et al*, 2025); however, the more distant *Zaprionus* genus – which branched from the *D. immigrans* subgroup and is now nested within the paraphyletic *Drosophila* genus – also has an extensive history of HTT with the *D. melanogaster* clade (De Setta et al., 2009, 2011; Simão et al, 2018; van Lopik et al, 2023).

To counteract potentially deleterious effects of TEs on the host genome, various host defence mechanisms have evolved to silence these elements. In *Drosophila*, this is mainly driven by the PIWI-interacting RNA (piRNA) pathway, a small-RNA based system that mediates TE silencing at both the co-transcriptional and post-transcriptional levels (Czech *et al*, 2018; Ozata *et al*, 2018). piRNAs against newly invaded TEs are thought to arise when the transposon inserts antisense into a piRNA cluster (Rozhkov *et al*, 2010; van Lopik *et al*, 2023). Conversely, TEs carry their own tissue-specific regulatory identity: the LTR regions contain transcription factor binding sites (TFBS) which enable transcriptional activity specifically in the cell types in which the elements are active (Faure *et al*, 1996; McDonald *et al*, 1997; Ludwig, 2002; Alizada *et al*, 2025; Rivera *et al*, 2025).

Recently, there has been a burst in the availability of *Drosophila* genome assemblies due, in part, to the more widespread availability of long read sequencing technologies (Kim *et al*, 2021, 2024). In this study, we took advantage of these large resources and annotated *Gypsy*-family TEs in 248 non-*D. melanogaster* drosophilid species. Our LTR-specific pipeline allowed us to capture and categorise more TEs than had been seen by previous methods. Using this resource of curated TEs, we focused on the proteins associated with cell-to-cell infectivity, ENV and sORF2. We show that sORF2 is highly conserved across drosophilids, whereas ENV is repeatedly lost. Lastly, we identified TFBS motifs associated with ENV presence and absence, allowing us to hypothesise that TEs hijack ovarian development pathways to ensure their transcriptional activation via host-intrinsic pathways. Altogether, our work provides unprecedented insights into the lifestyle of *Gypsy* TEs in drosophilids.

## RESULTS

### Most *Gypsy* TEs are present beyond the *melanogaster* clade

The *D. melanogaster* genome contains thousands of TE insertions (Mérel *et al*, 2020) representing over 120 distinct TE families, each defined by its own consensus sequence. A subset of these, belonging to the *Gypsy* family, are further divided into *Gypsy/gypsy*, some of which contain the *env* gene, *Gypsy/mdg1*, which harbour sORF2, and others (*mdg3, Osvaldo, chimpo*) (Senti *et al*, 2025). To understand the evolutionary origin of ENV and sORF2 and their role in defining TE lifestyles, we analysed 450 publicly available, long-read genome assemblies (Kim *et al*, 2024) representing 313 different species and spanning ten genera: *Amiota, Chymomyza, Caconexus, Drosophila, Hirtodrosophila, Lordiphosa, Scaptodrosophila, Scaptomyza, Zaprionus and Zygothrica*. We excluded 202 assemblies due to redundancy or lower quality (**Figure S1A**), leaving 249 species with a high-quality genome. Together, this collection spans approximately 50 million years of evolution (Suvorov *et al*, 2022) – with the lowest distance between species being 0.6 million years (between *D. cyrtoloma* and *D. melanocephala*) (Bonacum *et al*, 2005; Kumar *et al*, 2022) – as well as diverse geographic spread (**Figure S1B, C**).

We assessed the repeat landscapes of the 249 genomes using the Extensive *de novo* TE Annotator (EDTA) (Ou *et al*, 2019). Overall, we detected transposon content up to 55.5% *(L. clarofinis*) of the total genome size (**Table S1**). As expected, larger genomes were characterised by higher TE proportions (Pearson’s R = 0.94, p < 2.2×10^-16^) (**Figure S2A**), which may in part reflect more complete genome assemblies. Notable exceptions were the *Amiota, Chymomyza* and *Caconexus* genera, characterised by a lower-than-expected retroelement content, while their DNA transposon content followed the same trend as other species (**Figure S2B-D**). These three genera therefore have a different TE landscape compared with other drosophilids. An overview of genomic TE content across all 249 genomes is shown in **Figure 1A**. Overall, retroelements occupied a median 2.7-fold greater fraction of the genomes as compared with DNA transposons (Wilcoxon signed-rank test, p < 2.2×10^-16^) (**Figures 1A, D, S3**). Within the retroelements, LTR retrotransposons occupied a median 1.8-fold greater fraction than LINE elements (Wilcoxon signed-rank test, p < 2.2×10^-16^) (**Figures 1A, E, S3**). Lastly, among LTR retrotransposons, the *Gypsy* family has the highest genomic coverage (**Figures 1A, F, S3**), exceeding that of both *Copia* (median 25-fold lower, Wilcoxon signed-rank test, p < 2.2×10^-16^) and *Pao* (median 5-fold lower, Wilcoxon signed-rank test, p < 2.2×10^-16^). The *D. melanogaster* clade (i.e., ten species most closely related to *D. melanogaster*), ranging from *D. yakuba to D. melanogaster* (**Figure 1B**), all showed similar TE profiles, however, there were several differences in more distantly related species, for instance with higher overall TE content (39.68%) in *D. suzukii*. Lastly, since *Gypsy*-family TEs covered the largest fraction of drosophilid genomes, we asked how similar these were to the *D. melanogaster* reference TEs. To quantify this, we searched using RepeatMasker (Tarailo-Graovac & Chen, 2009) for sequences resembling the *D. melanogaster Gypsy*-family reference TEs (hereafter referred to *D. mel*-like) in each of the 248 genomes as well as the *D. mel* genome. Notably, we found species with up to 100% of their *Gypsy* TE content being *D. mel*-like (**Figure 1A, G, S3**). Species closely-related to *D. melanogaster* (**Figure 1B**), in the *Zaprionus* (**Figure 1C**) genus, and a few other species evolutionarily distant to *D. melanogaster* had the highest proportions of *D. mel*-like *Gypsy* TEs. We concluded that *D. melanogaster*’s TE content, measured in genomic coverage, is highly consistent across the *melanogaster* clade (**Figure 1A-B, S3**) and also comparable to *D. melanogaster* in more evolutionarily distant species, albeit with variation across different clades (**Figure 1A-G, S3**).

**Figure 1:**
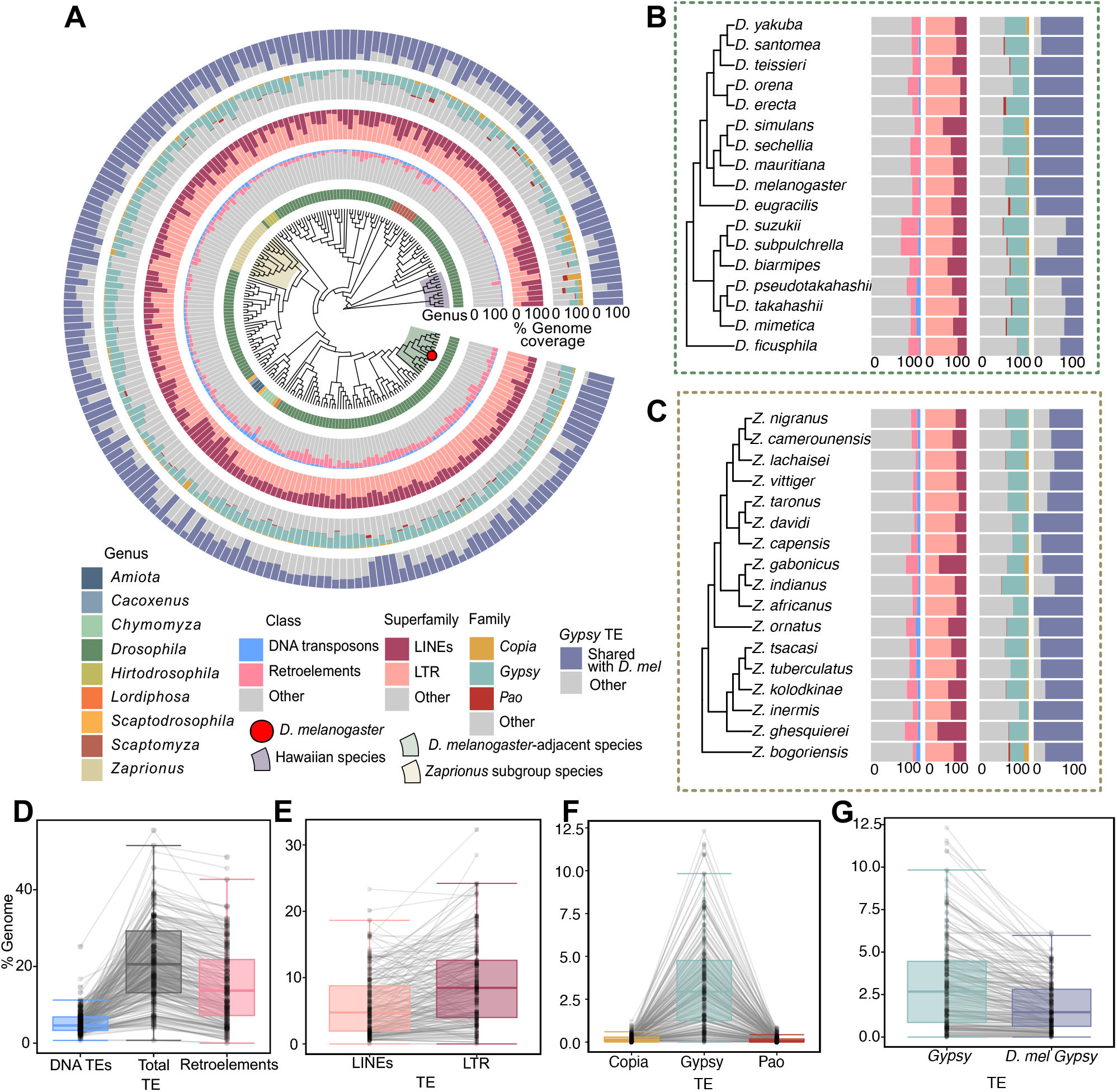
An overview of TE profiles in 249 drosophilid species. **A**: Phylogenetic tree of 249 drosophilid species. Genus is indicated by the innermost circle, followed by the percentages of TE class, retrotransposon superfamily and LTR family for each species as well as the percentage of *D. mel*-like *Gypsy*-family TEs, obtained by EDTA and RepeatMasker. Shaded regions on the tree highlight the *melanogaster* clade (green; *D melanogaster* shown as a red circle) and the *Zaprionus* genus (yellow), which are also shown in panel B and C. **B**-**C**: Phylogenetic trees show a zoom-in on the information shown in (A) for the *D. melanogaster* clade (B) and the *Zaprionus* genus (C). **D-G:** Paired box plots show the percentage of the genome covered by TEs, grouped by TE class (D), retrotransposon superfamily (E) and LTR family (F) and *D. mel*-like *Gypsy-*family TEs (G) for each species. Individual points between box plots are joined when they represent values from the same species.

### A pipeline to annotate *Gypsy*-family LTR retrotransposons in drosophilids

*Gypsy*-family TEs are the most diverse and prominent class in *D. melanogaster*. Having established similarities between the TE contents of *D. melanogaster* and other drosophilids, we focused on identifying which *D. melanogaster*-like TEs were present in the other species. This is challenging, as prevalent HTT combined with rapid evolution makes ortholog assignment across TEs particularly difficult. To this end, we used hidden Markov models (HMM) to identify *Gypsy*-family TEs in drosophilid genomes. We started with previously constructed and partially curated TE libraries (van Lopik *et al*, 2023) from which we filtered for *Gypsy*-family transposons and extracted their *POL* sequences. The *POL* sequences were used to construct a HMM profile of *POL*, which is highly conserved and has previously been used for *Gypsy* TE phylogenetic studies (Edlefsen & Liu, 2010; Senti *et al*, 2025; Voichek *et al*, 2025). The refined *POL* HMM profile was more generalisable across species and TE families compared with raw sequences and was used to identify TE insertions with high phylogenetic relatedness in each genome. Orthologs between the *D. melanogaster* consensus TEs and each species’ putative TEs were identified using reciprocal best hits. Briefly, we required each putative TE to be the best match for a *D. melanogaster* query TE and, conversely, for that *D. melanogaster* TE to be the best match for the putative TE sequence. Following this identification and assignment, downstream refining steps were applied to extend each hit into a full-length and high-quality consensus sequence (**Figure S4**). The pipeline outputs the consensus sequence of each available *D. melanogaster*-like *Gypsy*-family TE for each species (**Figure 2A-F**). When comparing to a previous study (Pianezza & Kofler, 2025), our framework captures a broader range of *D. melanogaster*-like TEs for most transposons tested (**Figure S5**).

**Figure 2:**
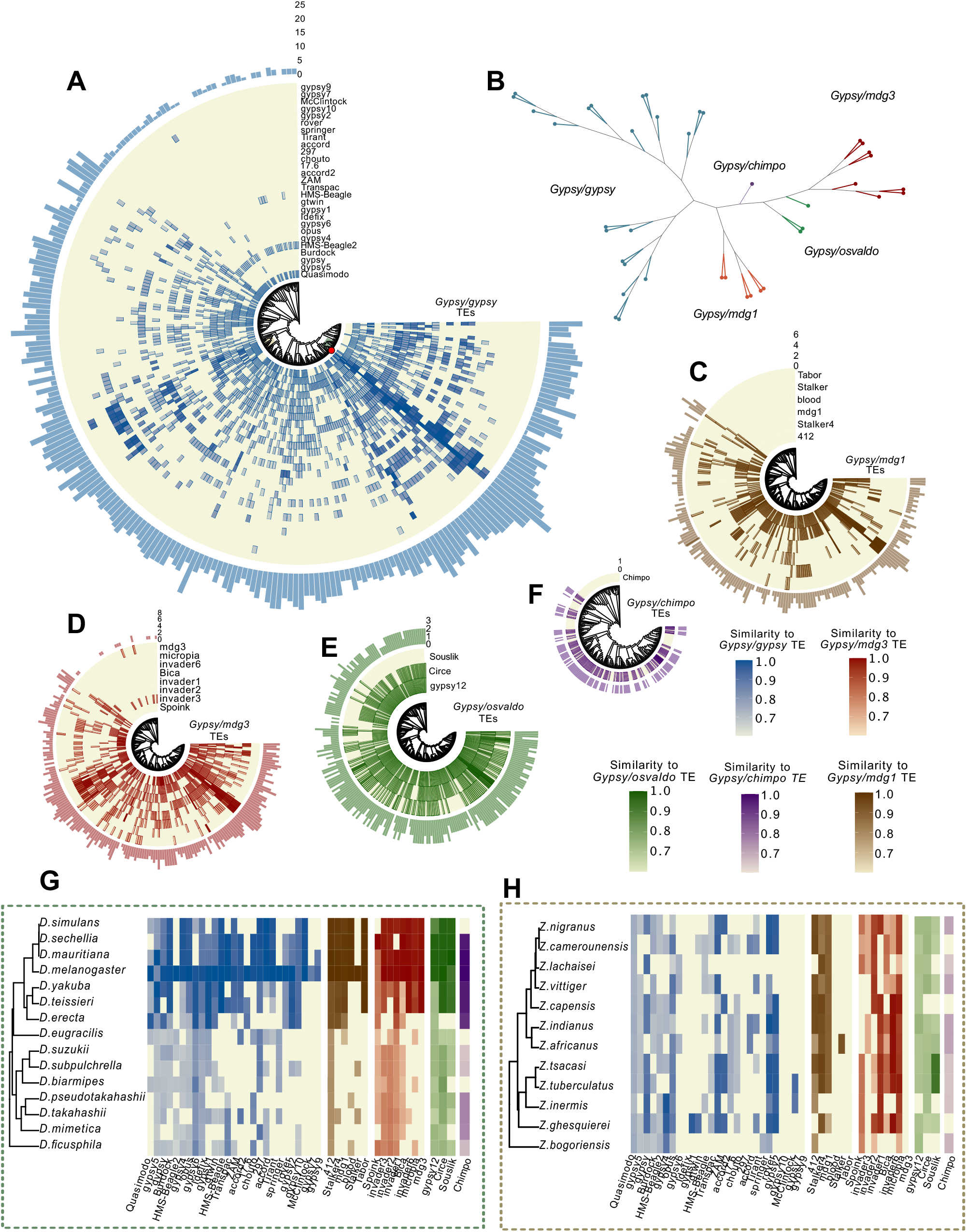
The presence of *Gypsy*-family LTR retrotransposons. **A**: A phylogenetic tree and accompanying heatmap show *Gypsy/*gypsy TE presence (≥70% similarity) and absence across 249 species. Heatmap colours indicate the similarity to the corresponding *POL* ORF in *D. melanogaster*. The bar plot represents the total number of *D. melanogaster Gypsy/*gypsy TEs identified in each species. **B**: Displayed is an unrooted phylogenetic tree of the D. *melanogaster Gypsy*-TE *POL* consensus sequences, coloured by the TE clade to which they belong. All internal nodes have a bootstrap support of ≥ 67%. **C-F**: Same as (A), but showing *Gypsy/mdg1* TEs (C), *Gypsy/mdg3* TEs (D), *Gypsy/osvaldo* TEs and *Gypsy/chimpo* TEs (F). **G-H**: Phylogenetic trees show in detail the same information as (A, C-F) for the *D. melanogaster* clade (G) and the *Zaprionus* genus (H).

In order to validate our pipeline, we sought to confirm that the sequences captured and annotated as Gypsy-family TEs were *bona fide* full-length elements rather than artefacts. LTR sequences provide a good estimate of this; the 5’ and 3’ LTRs of an LTR retrotransposon are identical at the time an insertion takes place, a by-product of the reverse transcription mechanism (Hughes, 2015). Similar 5’ and 3’ LTR sequences therefore suggest a recent insertion. Here, we measured this by annotating the LTRs of the identified TE consensus sequences. We found that, for all TEs found in more than five species, 78% of LTR pairs in the consensus sequences shared >95% identity (**Figure S6**), suggesting that the TEs identified do, indeed, come from insertion events.

To validate our annotations further, we tested whether they are potentially controlled by the piRNA pathway, through mapping small RNA sequencing (sRNA-seq) data, where available, to our generated *Gypsy* TE libraries. As the piRNA pathway silences TEs by sequence complementarity, we expect piRNAs, captured by sRNA-seq, to map antisense to these sequences. By analysing published sRNA-seq data (Chirn *et al*, 2015; Lewis *et al*, 2017; Mohammed *et al*, 2018; Kotov *et al*, 2019; Vedanayagam *et al*, 2021; Zhao *et al*, 2021; van Lopik *et al*, 2023; Son *et al*, 2025), we first confirmed that piRNAs sequenced in each species map to the TE library we constructed from that species (**Figure S7**, diagonal boxes). Next, we confirmed that piRNAs from the *melanogaster* clade and *Zaprionus* genus map among each other’s TEs (**Figure S7**), as would be expected for species with many similar TEs. Moreover, we showed that the piRNAs mapping to these TEs show the expected nucleotide frequencies (5’ U bias) (Brennecke *et al*, 2007), lengths (23-29 nt) and antisense mapping (**Figure S8**). In conclusion, piRNA sequencing data therefore aligns with the idea that our TE libraries largely consist of active or formerly active TEs, which are under the control of the piRNA pathway. Therefore, our pipeline provides a sensitive and accurate way of detecting high-fidelity *Gypsy* TE insertions and constructing consensus sequences of individual TE families in distantly related species.

### *Gypsy-*family LTR retrotransposons are widespread across drosophilids

Having established a resource containing a total of 3,438 Gypsy-family consensus sequences across drosophilids, we were now able to study their evolutionary dynamics. Strikingly, conservation of *Gypsy*-family TEs varied widely across the phylogenetic tree, with anywhere between none or all (in *D. melanogaster*) of the 43 tested TEs – spanning five TE families (**Figure 2B**) – found in individual species (at ≥70% similarity). *Gypsy/gypsy* (**Figures 2A, S9A**), *Gypsy/mdg3* (**Figures 2D, S9A**), and *Gypsy/osvaldo* (**Figures 2E, S9A**) were found across the drosophilid phylogenetic tree, whereas *Gypsy/mdg1* (**Figures 2C, S9A**) and *Gypsy/chimpo* (**Figures 2F, S9A**) elements were not found in the most distantly related species. As expected, *D. melanogaster* TEs were most frequently found, and with highest similarity, in closely related species (*D. simulans-D. melanogaster*) (**Figure 2G**).

Moreover, the *Zaprionus* genus, known for extensive HTT with *D. melanogaster* (De Setta et al., 2009, 2011), shows high similarity between POL proteins for some *Gypsy* TEs (*ZAM, rover*) (**Figure 2H, S9B-C**). Reassuringly, an alignment and resulting tree created from all *Gypsy* TE *POL* sequences shows that *POL* sequences annotated as the same *D. melanogaster Gypsy* clade cluster together (**Figure S10A**), reflecting their distinct evolutionary trajectories. Moreover, individual *POL* consensus sequences cluster together by TE and irrespective of species (**Figure S10B-G**), consistent with a single origin of each TE. In conclusion, we detect widespread and conserved *Gypsy* TEs in a range of *Drosophila* species.

The *POL* sequence similarity between our annotated *Gypsy* TEs in species and their counterparts in *D. melanogaster* varied between 70% (the cutoff of our pipeline) and 100% similarity (**Figures 2A-H, S9**). This suggests that TE ORFs may evolve between species. To better study the selective pressures and mutational biases that shape the evolutionary dynamics of the *Gypsy*-family TEs in drosophilids, we calculated the non-synonymous to synonymous mutation (*dN/dS*) as well as the transition to transversion (*Ts/Tv)* ratios of our annotated *POL* sequences. The *dN/dS* and *Ts/Tv* scores show a high degree of purifying selection of the ORFs, with all *dN/dS* scores below 1 (**Figure 3A-B**). POL has *dN/dS* values on average 0.097±0.005, in the same range as other *Drosophila* protein-coding genes (Domazet-Loso & Tautz, 2003). A likelihood ratio test (LRT) also favoured the estimated selection models over neutral (*dN/dS*=1) evolution (LRT statistic range: 71.7–34,282, p<10^-15^ for all tests). The higher *dN/dS* and *Ts/Tv* values for ENV compared to other ORFs suggest that the sORF2 and ENV ORFs may be undergoing purifying selection, like POL, albeit to a lesser degree (**Figure 3A-B**). These observations suggest that *Gypsy*-family elements in drosophilids are subject to purifying selection, that they remain functionally active, and are under evolutionary pressure to maintain the functionality of their coding sequences. There is, however, heterogeneity in these evolutionary constraints, implying that some ORFs are more strictly regulated than others.

**Figure 3:**
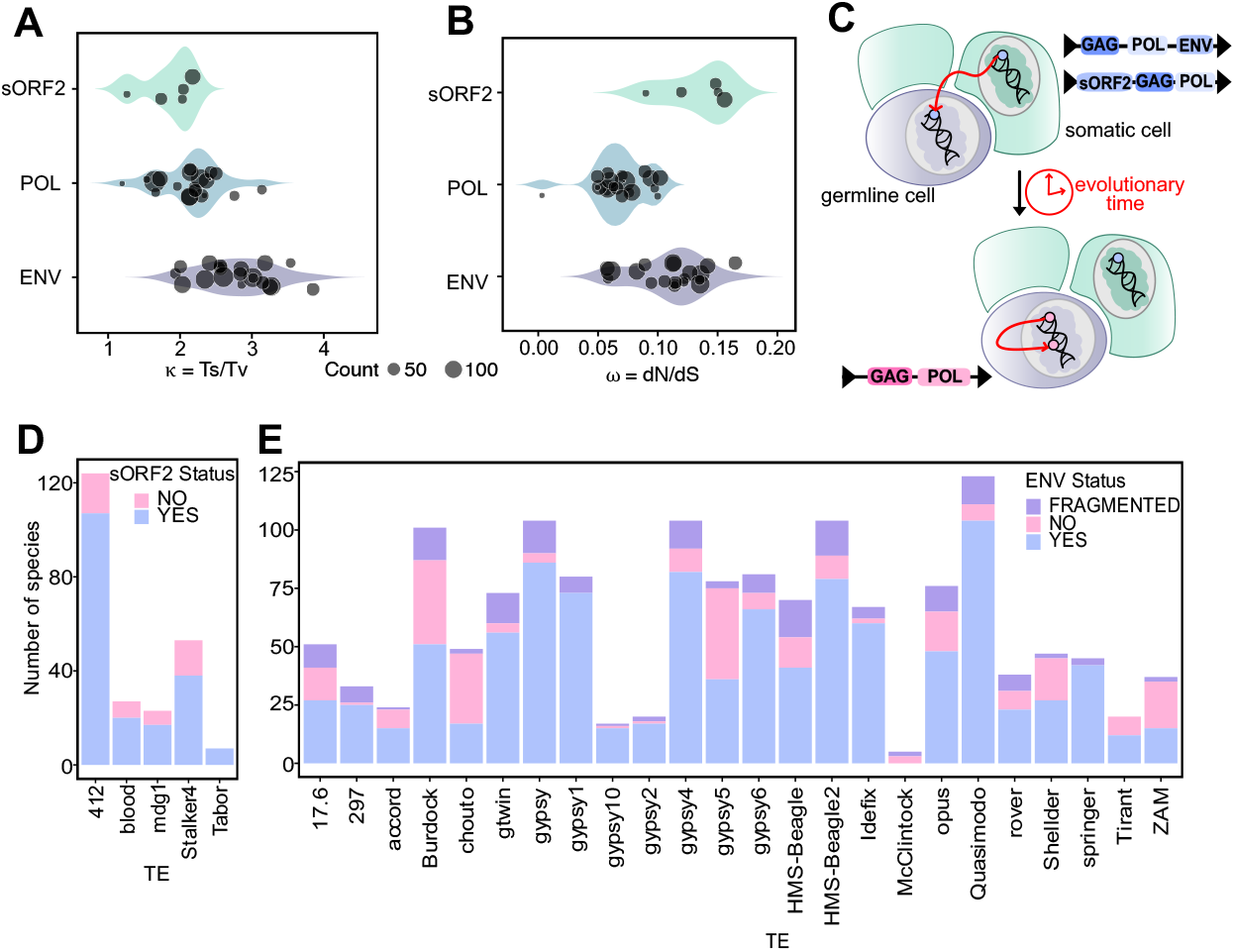
Analysis of ENV and sORF2 indicates complementary functions and contrasting patterns of evolution. **A**: Violin plots show the distribution of *Ts/Tv* values for ORFs in *Gypsy*-family TEs. **B**: Shown is the same as A, but *dN*/*dS* values are displayed. **C**: A schematic diagram shows the current model for *ENV* loss. TEs are initially active in the soma and require specialised ORFs to infect the germline. Upon a shift from somatic to germline expression, the ORF becomes redundant and, through natural drift, the ORF becomes lost or inactive. **D**: A bar plot shows the status of the *sORF2* ORF in TE consensus sequences by number of species. **E**: A bar plot shows the status of the *ENV* ORF in TE consensus sequences by number of species. *ENV* “presence” was considered to be an identified ENV equal to or longer than 900 nucleotides, while “fragmented” ENV was an ENV detected shorter than 900 nucleotides.

### Gypsy TEs show distinct insertion preferences

To better understand the dynamics of past TE invasions, we next examined the number and location of insertions for each TE subfamily. Many insertions were found to be highly similar to the consensus sequence (**Figure S11**), indicating that they may have recently invaded a species, while others were more diverged, suggesting older TE invasions which have allowed for the accumulation of mutations. For instance, *297* is likely younger in *D. melanogaster* (0.078% mean divergence across insertions) than in *D. mauritania* (2.2% mean divergence across insertions). Interestingly, the same TE can have different divergence profiles across species, indicating potentially differentially successful invasions. Similarly, several TEs have successfully invaded (i.e., have a large number of full-length copies) different species (**Figure S12**), and most species only have one such recently invading TE. Although TEs insert largely randomly in the genome, some TEs may have preferential insertion sites. Although TE divergence profiles varied across species (**Figures S13, S14**), we did not observe a clear correlation between insertion location and TE divergence. Some TEs such as *blood* did, however, have more insertions in intragenic and promoter regions than others (**Figure S15**).

### Phylogenetic incongruences are prominent in the evolutionary history of *Gypsy*-family TEs

The evolutionary history of TEs is known to be complex and to contain many deviations from vertical inheritance, including HTT. Large incongruences between *POL* sequence phylogenetic trees and the species trees (**Figure S16**) highlight multiple putative instances of HTT in the evolutionary history of all *Gypsy/gypsy* TE families. To quantify all potential instances of HTT among the *Gypsy* TE phylogenetic tree, we identified species which had swapped “neighbours” between the species and TE trees. These phylogenetic incongruences exist both between closely related species, which may in part be due to incomplete lineage sorting or hybridisation, and across clades, indicative of HTT, with multiple such incongruences observed between the melanogaster group and *Zaprionus* genus (**Figure S17A**), as recently reported (Pianezza & Kofler, 2025). We speculate that the genomes with few *D. mel*-like TEs may be explained by limited opportunities for transposon exchange due to geographic segregation. The Hawaiian species (*D. silvestris, D. tanythryx*) are good examples of this as they consistently only share four TEs with *D. melanogaster* (*Circe, gypsy12, Quasimodo, HMS-Beagle2*) (**Figure S9A**). Indeed, TEs seem to have been passed to the Hawaiian species through a single putative HTT event (**Figure S17B**). Tropic niche segregation (Markow & O’Grady, 2008), for instance with cactophilic species (*D. mojavensis*), may contribute to the same phenomenon and has previously been shown to shape species’ HTT landscapes (Carvalho *et al*, 2023).

### *Gypsy/gypsy* and *Gypsy/mdg1* TEs have contrasting patterns of evolution

To date, the *Gypsy/gypsy* and *Gypsy/mdg1* families are the only *Drosophila Gypsy* TEs known to contain proteins which enable cell-to-cell infectivity (Senti *et al*, 2025; Voichek *et al*, 2025) (**Figure 3C**). Moreover, the *ENV* protein is ancestral to *Gypsy*/*gypsy* and has been lost across many *D. melanogaster* TEs (Senti *et al*, 2025). A change of expression of a TE from the soma to the germline would likely be accompanied by the gradual loss of the ENV protein due to random drift. To study this in detail, we focused on the conservation of *sORF2* and *ENV* across TE insertions in drosophilids. We asked whether full-length *ENV* and *sORF2* ORFs were consistently present or absent for a given *Gypsy/gypsy* or *Gypsy/mdg1* TE consensus sequence across all species which it has invaded.

The sORF2 protein was consistently present in ≥80% all *Gypsy/mdg1* TE consensus sequences (**Figure 3D**). The consistently high prevalence of sORF2 in *Gypsy/mdg1* TEs, the presence of purifying selection (**Figure 3A-B**), combined with the broad prevalence of these TEs across species (most notably the TE *412*, which is found in 124 out of 249 species, suggests that sORF2 is a near-universal component of *Gypsy/mdg1* TEs. In contrast, the ENV protein is not stably present or absent across *Gypsy/gypsy* TEs (**Figure 3E**). Rather, it shows a high rate of ENV loss events (16 TEs with <80% ENV presence) throughout TE evolution. Moreover, ENV can also be detectable but fragmented and thereby non-functional, suggesting that it is no longer active but may have been in the past, as previously suggested in *D. melanogaster (Senti et al, 2025)*. Interestingly, ENV presence/absence is not concordant with the species or TE trees (**Figure S16**), suggesting multiple, independent, gain or loss events across its evolutionary history. Notably, there were no co-occurrences of the sORF2 and ENV proteins, suggesting that they provide complementary and distinct mechanisms to enable a somatic niche for a TE.

### *Gypsy/gypsy* TE expression patterns correlate with specific TFBSs

A TE must contain all the regulatory elements required for its expression. The LTR regions of TEs contain small promoter regions that determine their transcription in specific tissues and cell types. They are known to contain TFBSs which determine TE expression patterns and levels (**Figure 4A**). For instance, the transcription factor *traffic jam (tj)*, in addition to its role in ovarian somatic cell development (Li *et al*, 2003; Kawashima *et al*, 2003), activates TEs as well as piRNA pathway components which, in turn, repress TEs (Alizada *et al*, 2025; Rivera *et al*, 2025). To understand whether additional transcription factors may regulate transposon expression, we performed a computational screen for TFBS motifs in TEs in drosophilids. To focus the screen, we included only motifs for transcription factors expressed in the *D. melanogaster* ovary (**Figure 4B**). We asked whether the presence or absence of motifs in the LTR region of a TE consensus sequence could be used to predict its ENV status. This relies on the hypothesis that ENV-containing TEs are expressed in the soma and TEs which do not contain functional ENV are expressed in the germline.

**Figure 4:**
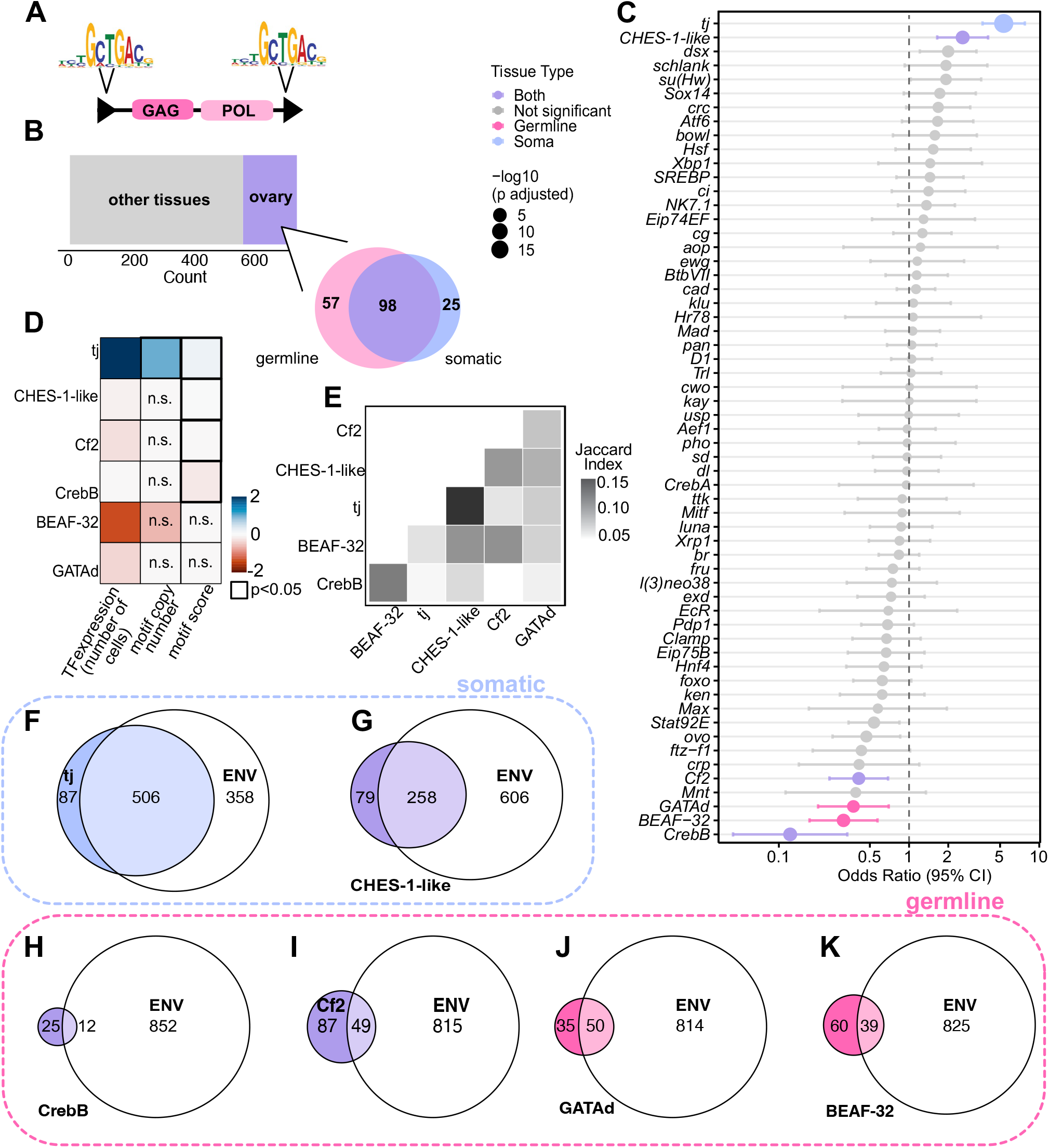
Specific TFBSs are associated with ENV presence or absence. **A**: A schematic diagram shows the structure of an LTR retrotransposon and TFBS presence in the LTR regions. **B**: A bar plot shows the number of ovary-expressed and non-ovary-expressed transcription factors in *D. melanogaster* and a Venn diagram shows the distribution of somatic, germline, and whole-ovary-expressed transcription factors in *D. melanogaster* ovaries. **C**: The ORs for associations between TFBSs predicted in TE LTRs and *ENV* presence is displayed. The transcription factors shown are highly expressed in ovarian cells, defined as being detectable through scRNA-seq (Li *et al*, 2022) in at least 20% of germline, somatic cells, or both. **D**: A heatmap shows the log_2_ fold changes of the medians for the indicated metrics (proportion of cells with detectable expression, per-species number of TFBS motifs, or score of the motifs) for the six identified transcription factors between TEs containing full-length *ENV* and TEs which do not contain full-length *ENV*. The associated *p*-values, where significant, are the result of a Wilcoxon rank-sum test and are displayed in the heatmap. Crosses indicate non-significant ratios. **E**: A heatmap shows the Jaccard index for TFBS co-occurrence in TE LTR regions for the six transcription factors with significant ORs. **F-K**: Venn diagrams show the overlap between TEs containing full-length *ENV* and TEs containing the TFBS motif in their LTR regions, for the six transcription factors found to be significantly associated with *ENV* presence or absence: *tj* (F), *CHES-1-like* (G), *CrebB* (H), *Cf2* (I), *GATAd* (J), and *BEAF-32* (K).

We used a total of 1,267 5’ LTR region sequences of which 864 were associated with a full-length ENV ORF. TE families where all species had the same ENV status (e.g., absent in all *McClintock* TEs) were excluded from this analysis. Using a mixed effects logistic regression with TE family modelled as a random effect, we identified that *tj* (Odds Ratio [OR] = 5.06, *p*_adj_ = 5 × 10^-20^) and *CHES-1-like* (OR = 2.78, *p*_adj_ = 8 × 10^-7^), both of which are somatically expressed, were significantly associated with ENV presence. Moreover, four transcription factors expressed in the germline were significantly associated with ENV absence: *Cf2* (OR = 0.323, *p*_adj_ = 4 × 10^-6^), *GATAd* (OR = 0.549, *p*_adj_ = 0.043), *CrebB* (OR = 0.113, *p*_adj_ = 1 × 10^-5^) and *BEAF-32* (OR = 0.340, *p*_adj_ = 1 × 10^-4^) (**Figure 4C**).

Of note, while three of six identified transcription factors were detectible in both the germline and the soma (**Figure 4C**, purple hits), they were in all cases most broadly detectible in the tissue where they were predicted to regulate TEs (**Figure 4D**). Comparing TFBS motifs across TEs with our without ENV, we found that motif scores were, for *tj, CHES-1-like, Cf2* and *CrebB*, significantly stronger in the TEs they were predicted to regulate (**Figure 4D, S18**), and *tj* contained significantly more TFBSs in the LTRs of TEs without an ENV protein (**Figures 4D, S19**). The predictors of somatic expression, *tj* and *CHES-1-like*, were also the most likely to co-occur as the pair had the highest Jaccard index (**Figure 4E**). *tj*’s binding motif was the most abundant and present in at least one species for nearly all the *Gypsy/gypsy* TEs tested (**Figure 4F**). Moreover, TFBSs associated with ENV presence were primarily found in TEs which contain ENV (**Figures 4F, G**) and, conversely, ones associated with ENV absence were primarily found in TEs which lack the ORF (**Figure 4H-K**).

Finally, conservation analysis of our most significant hits confirmed that they are present and highly conserved across drosophilid species, further indicating their potentially conserved role in determining TE expression patterns (**Figure S20**).

## DISCUSSION

Here, we took advantage of recent efforts in long-read whole-genome sequencing across drosophilids (Kim *et al*, 2021, 2024) to study TEs at a large scale and across multiple species. Previously, similar genome assemblies allowed the identification of piRNA clusters in multiple non-*D. melanogaster Drosophila* species, yielding insights into the regulation of TEs (van Lopik *et al*, 2023), as well as enabling evolutionary insights into other repetitive genomic elements (Gebert *et al*, 2025). High-quality genome assemblies have also enabled recent in-depth studies of coevolution of TEs with their host, such as in *D. virilis* (Rezvykh *et al*, 2026). Here, we instead used the available drosophilid genomes to construct *Gypsy*-family TE libraries for 249 species. The resulting resource allowed extensive evolutionary studies on TEs across multiple species spanning 50 million years of evolution.

Our results highlight the widespread nature of TE transfer, as putative HTT events were observed in the evolutionary history of all *Gypsy*-family TEs studied. In line with similar recent findings in other organisms, this establishes TE mobility, including HTT and introgression as key drivers in shaping the mobile genome (Romeijn *et al*, 2025) and particularly in *Drosophila* (Jordan *et al*, 1999; Scarpa *et al*, 2025). In drosophilids, recent work has characterised potential HTT events among species and suggested that HTT events occur when species are in close geographical proximity (Pianezza & Kofler, 2025). Currently, the vast number of phylogenetic incongruences between TE trees and host species trees make it difficult to pinpoint an exact donor and acceptor of putative HTT events. While clustering and topological incongruence between the TE and species phylogenetic trees is a good indicator of HTT (Peccoud *et al*, 2017; Baidouri *et al*, 2014), several alternative methods exist to identify HTT events. These include pairwise comparisons of evolutionary divergence of all shared genes followed by comparing the TE divergences against a baseline determined by those of BUSCO genes (Pianezza & Kofler, 2025). Moreover, the vertical hybridisation and inference of congruence analysis (VHICA) method compares signatures of selection (*dS*) between genes of interest and a user-determined baseline of conserved, vertically transferred genes (Wallau *et al*, 2016). Despite this, HTT is difficult to differentiate from other phenomena such as recombination and coalescence. We hope that our TE annotations combined with larger-scale genomic surveys and future tree reconciliation methods will enable this.

TE-encoded proteins of viral origin that enable cell-to-cell movement of the elements are of interest due to their ability to enable a TE to infect the germline (Senti *et al*, 2025; Voichek *et al*, 2025). Previous work has indicated that insertions of the same TE, *rover*, in *D. melanogaster* can encode both active and inactive protein variants, a distinction reflected in the insertions’ expression patterns (Senti *et al*, 2025). On a genus-wide level, our data show that *Gypsy/mdg1* TEs have a highly conserved sORF2 protein, present in nearly all detected consensus sequences across drosophilids, whereas *Gypsy/gypsy* TEs have lost the ENV protein repeatedly throughout their evolutionary histories. A central premise guiding our analyses is the well-supported dichotomy in *D. melanogaster* that TEs containing ENV or sORF2 are expressed in the soma whereas elements lacking a functional copy of either of these ORFs reside in the germline (Senti *et al*, 2025; Voichek *et al*, 2025). Although we did not directly assay tissue-specific expression for the drosophilid species analysed in this study, we consider this assumption highly robust due to the extensive evidence in *D. melanogaster* and known role of ENV and FAST-like proteins in mediating cell-to-cell transfer. Consistent with its evolutionary plasticity, *ENV* has a higher, on average, *dN*/*dS* ratio relative to the other proteins and independently of the species or TE phylogenies.

Transcription of TEs is mediated by TFBSs in their LTR regions (Alizada *et al*, 2025; Rivera *et al*, 2025; Milyaeva *et al*, 2023). In *Gypsy/gypsy* TEs, we identified two transcription factors significantly associated with somatic expression, *tj* and *CHES-1-like*, and four significantly associated with germline expression: *Cf2, GATAd, CrebB* and *BEAF-32*. These factors were previously characterised as crucial in *D. melanogaster* gonad development (Yu *et al*, 2016; Hsu *et al*, 2001; Chen *et al*, 2022; Kawashima *et al*, 2003; Li *et al*, 2003). Interestingly, BEAF-32 also regulates the non-LTR TEs and the piRNA pathway (Sokolova *et al*, 2023). Hijacking key developmental pathways likely means that the transcription factors required for TE expression will be present following a jump to another species. Another benefit is likely that the host is unable to render a key transcription factor dysfunctional to halt a TE invasion. Some transcription factors whose TFBSs motifs were not significant hits in our screen may still be functional in activating TE expression and their redundancy may serve as a fail-safe mechanism to ensure transcription of the TE insertion across many cellular niches.

In conclusion, we present a resource containing *D. melanogaster* TE consensus sequences in 249 high-quality drosophilid genomes. We highlight the distinct evolutionary mechanisms by which some ORFs evolve and shed light on the highly conserved mechanisms TEs use to ensure their transcription in their host genomes. Our results and dataset will enable studies of TEs co-evolution with the host genome in the future, particularly focusing on TE transcription and cell type-specific expression.

## MATERIALS AND METHODS

### Genome assemblies

We used previously published resources of long-read assemblies (Miller *et al*, 2018; Kim *et al*, 2021, 2024) as listed in **Table S3**.

### Genome assembly quality control

Genome quality control was carried out using calN50 (https://github.com/lh3/calN50), checking the number of sequences (NN) and genome contiguity (N50). A genome assembly was considered to have passed the quality threshold if N50>1,000,000 and NN<3,000. For species with more than one available genome assembly, the highest quality assembly was kept.

### De novo TE annotations

Initial *de novo* TE libraries were built for each genome using RepeatModeler (v2.0.7) (Flynn *et al*, 2020) *BuildDatabase* followed by *RepeatModeler* (-threads 4 -LTRStruct). These were inputted as the RepeatModeler library (--rmlib) into EDTA (v2.2.2) (Ou *et al*, 2019) (--species others --step all --anno 1 --threads 4 --force 1). SINE elements were not included in this analysis due to low genomic coverage, likely due to difficulties being captured by the automated tools used (Kapitonov & Jurka, 2003; Nefedova & Kim, 2009). The percentages of the genome covered by each TE class, superfamily and family were extracted from the RepeatMasker *.tbl output file generated by EDTA. To quantify how much of the *Gypsy*-family coverage that was *D. mel*-like, we compared the total *Gypsy*-family genomic coverage to that of the *D. melanogaster* reference transposon library estimated using RepeatMasker (Tarailo-Graovac & Chen, 2009). To compare the genomic proportions of TE groups, a Wilcoxon signed-rank test was utilised. We opted for a paired non-parametric test as the percentages compared are paired by host species. We used the wilcox.test function from the stats (v4.5.1) package in R (v4.5.1) with the paired=TRUE flag.

### Phylogenetic tree visualisation

A previously generated phylogenetic tree for 704 drosophilidae species (Finet *et al*, 2021) was used to display our species onto a phylogenetic tree. The phylogenetic tree was imported using treeio (v1.28.0) (Wang *et al*, 2020) and species not included in our study were removed. The tree and metadata were visualised using ggtree (v3.12.0) (Xu *et al*, 2022) and ggtreeExtra (v1.19.1) (Xu *et al*, 2021).

### ORF identification using HMMs

TE libraries (van Lopik *et al*, 2023) from 193 previously studied *Drosophila* genome assemblies were filtered to only include *Gypsy*-family LTR retrotransposons. Their *POL* sequences were extracted by mapping all consensus sequences to RepeatPeps.lib in RepeatMasker (v4.1.2) (Tarailo-Graovac & Chen, 2009) using blastx (v2.16.0) (Camacho *et al*, 2009). *GAG* and *ENV* ORFs were extracted in the same way. The resulting ORF nucleotide sequences were assembled into a HMM using hmmer’s (v3.4) hmmbuild (Eddy, 2008, 2009, 2011) using Henikoff position-based weights. A HMM for *sORF2* was constructed using available *sORF2* sequences (Voichek *et al*, 2025). The *POL* HMM was used to identify *POL* sequences in all drosophilid genomes using nhmmer with default parameters. True hits were considered to be ones with an e-value <10^-20^ and length >2,500 nucleotides. The HMMs developed are available on https://github.com/annamaria23/retrotransposon-atlas/tree/main/2_ORF_HMMs.

### TE annotation

The *D. melanogaster* reference *Gypsy* TE database was built using a pre-existing, manually semi-curated, TE database (https://github.com/bergmanlab/drosophila-transposons) to which we manually added eight recently identified *Gypsy*-family TEs: *chouto, Chimpo, Bica* (Senti *et al*, 2025), *MLE, Souslik, Transib1* (Pianezza *et al*, 2025), *Spoink and Shellder* (Scarpa *et al*, 2025). In cases where *POL* was not already annotated, it was identified using the same HMM method as for putative TEs (cf. *ORF identification using HMMs*) or using blastx (v2.16.0, default parameters) against *POL* sequences in RepeatPeps.lib (RepeatMasker, v4.1.2) and the longest sequence was selected and manually validated. MMseqs2 (v18.8cc5c) (Steinegger & Söding, 2017) *createdb* followed by *rbh* and *convertalis* with default parameters were used to identify reciprocal best hits between all putative *POL* sequences in a genome and the *POL* sequences of the curated *D. melanogaster* TEs.

### Consensus sequence creation

To create full-length TE consensus sequences, the annotated *POL* ORFs were mapped back to each genome using blastn (v2.16.0) (Camacho *et al*, 2009) filtering for 90% identity, 1e-20 E value cutoff, and for the resulting hits to be longer than half that of the query sequence. The top 20 sequences were selected, based on E value. The resulting sequences were extended 5,000 nucleotides up- and down-stream using bedtools slop (v2.31.1) (Quinlan & Hall, 2010) In cases where three or more sequences were present, a multiple sequence alignment (MSA) was constructed using mafft (v7.526, --reorder --auto) (Katoh *et al*, 2002; Katoh & Standley, 2013). CIAlign (v1.1.4) (Tumescheit *et al*, 2022) was used to refine the MSA by filtering divergent sequences (--remove_divergent --remove_divergent_minperc 0.3 --remove_short) and low-coverage regions (--crop_divergent --crop_divergent_min_prop_nongap 0.8 --crop_divergent_min_prop_ident 0.8 --crop_ends --remove_insertions --insertion_max_size 3000) and generate a consensus sequence of the TE (--make_consensus). The full consensus sequence of the putative TE was mapped back to the genome using blastn (v2.16.0) (Camacho *et al*, 2009) with default parameters and the 5’ and 3’ ends of the TE sequences were trimmed to remove nucleotides with low mapping coverage to the genome using a custom Perl (v5.32.1) script (crop_zero_coverage.pl from https://github.com/susbo/Drosophila_unistrand_clusters) (Bornelöv & JvanLopik, 2024). In cases where fewer than three *POL* sequences were identified in the genome from the initial blast step (this would represent TEs with only one or two insertions in the genome), the one with the lowest E value was selected and considered the consensus sequence.

### Annotation of TE consensus sequences

To obtain the LTR regions of putative TEs, blastn (v2.16.0) was used on the sequence against itself. The longest identified sequences where the 5’ LTR ended before the start codon of the *GAG* ORF and the 3’ LTR began after the stop codon of the *POL* ORF were considered to be the LTR regions. TE sequences were trimmed such that they began at the first nucleotide of the 5’ LTR and ended at the last nucleotide of the 3’ LTR.

### TE annotation benchmarking

We compared the annotations of this work with those of a recent study (Pianezza & Kofler, 2025) found on: https://github.com/rpianezza/Drosophilids-TE-biogeography/blob/main/D.melanogaster/data/TEs.clean.score. We used the provided metadata file (te-metadata.csv) to covert the TE names and filtered for a TE score ≥ 0.5, following the previously published methodology. Lastly, we filtered for *Gypsy-*family elements and only included TEs and species for our downstream analysis that were included in both studies.

### Similarity between LTR sequences

blastn (v2.16.0, default parameters) between the 5’ and 3’ LTRs of TE consensus sequences was used to obtain their pairwise similarity.

### Publicly available sRNA-seq data

We used publicly available sRNA-seq data from previous studies (Chirn *et al*, 2015; Lewis *et al*, 2017; Mohammed *et al*, 2018; Kotov *et al*, 2019; Vedanayagam *et al*, 2021; Zhao *et al*, 2021), which were also used in (van Lopik *et al*, 2023), to which we added sRNA-seq data from *D. triauraria* and five *Zaprionus* species (Son *et al*, 2025). An overview of the libraries used is available in **Table S2**.

### sRNA-seq

All sRNA-seq data were processed using the same analysis pipeline as (van Lopik *et al*, 2023). Briefly, after removing an abundant rRNA, sequencing adapters, any flanking random nucleotides and reads mapping to miRNAs, as described (van Lopik *et al*, 2023), the remaining reads were mapped to the TE consensus sequences using bowtie (v1.2.3, -S -n 2 -M 1 -p 20 --best --strata --nomaqround --chunkmbs 1024).

### *dN*/*dS* and *Ts*/*Tv* scores

The HMM-based consensus ORF sequence was refined and then translated using EMBOSS’ *getorf* (v6.6.0.0, -find 3 -minsize 900) (Rice *et al*, 2000) and *transeq* functions. After duplicate sequences were removed using seqkit rmdup, amino-acid MSAs were obtained using mafft (v7.526) (Katoh *et al*, 2002; Katoh & Standley, 2013) with default parameters. The amino-acid MSA and sequence files were used to generate a codon alignment using PAL2NAL (v14) (Suyama *et al*, 2006) and phylogenetic trees were generated from this MSA using IQTREE2 (v2.3.6) (Minh *et al*, 2020) with the -m MFP flag. *dN/dS* and *Ts/Tv* scores were obtained using PAML’s *codeml* (v4.10.7, CodonFreq2; model 0; NSsites 0; fix_omega 0; omega 1) (Álvarez-Carretero *et al*, 2023) function. The neutral evolution model (M(0)=0, fix_omega 1) was compared to the selection model (M(0)=1, fix_omega 0) using a likelihood ratio test (2* [lnL_M1_-lnL_M0_]) for each TE ORF consensus sequence individually. The resulting likelihood ratio test statistic was compared against a χ^2^ distribution with one degree of freedom.

### Identification of individual TE insertions and proximity to protein-coding genes

TE libraries for each genome were searched in the species’ genome using RepeatMasker (Tarailo-Graovac & Chen, 2009), specifying the library with the -lib flag, BUSCO (v6.6.0) for the *drosophilidae_odb12* lineage with default parameters was used to identify protein-coding genes in each genome. Lastly, bedtools (v2.31.1) (Quinlan & Hall, 2010) *closest* was used to identify the closest protein-coding gene to each TE insertion.

### Phylogenetic trees from POL TE consensus sequences

MSAs of *POL* nucleotide sequences were generated using mafft (v7.526) (Katoh *et al*, 2002; Katoh & Standley, 2013) with default parameters and refined using CIAlign (Tumescheit *et al*, 2022) (v1.1.4, --crop_divergent --crop_divergent_min_prop_nonga p 0.8 --crop_divergent_min_prop_ident 0.8 --remove_divergent --remove_divergent_minperc 0.3 --crop_ends --remove_insertions --insertion_max_size 3000 --remove_short). For individual TE *POL* ORFs, phylogenetic trees were constructed from the resulting MSAs using IQTREE2 (v2.3.6) (Minh *et al*, 2020), selecting the best substitution model using ModelFinderPlus (-m MFP). The tree search was initiated from a pool of 100 initial parsimony trees (--ninit 100). The reliability of the internal branches was tested using the SH approximate likelihood ratio test (-alrt 1000), approximate Bayes test (-abayes) and ultrafast bootstrap (-B 1000) with an additional optimisation step using Nearest-Neighbor Interchange applied to each bootstrap replicate (-bnni). A fixed random seed (--seed 42) was used. When computing phylogenetic trees for all TE ORFs, FastTree (v2.2.0) (Price *et al*, 2009) with default parameters and removed identical sequences.

### Identifying putative HTT events

Putative HTT events were called by comparing the phylogenetic topologies of the TE tree and species tree using the ete3 (v3.1.3) (Huerta-Cepas *et al*, 2016) package in Python (v3.10.8). For each species, its closest relatives are defined in both the species tree and the TE tree as the descendant leaves with the same parent node. For each species, if its set of “neighbour” species in the gene and species trees do not share at least one element, HTT is considered to have taken place between the target species and the “new” relative on the TE tree. To ensure reliability, only nodes with an ultrafast bootstrap support value above 70% were considered.

### Distance between *ENV*-containing TEs in phylogenetic trees

The *cophenetic()* function was used in R (v4.5.1) to calculate distances between selected nodes in trees. The mean value of the lower triangle of these distances was then taken.

### Motif discovery

Motifs in LTR regions were identified from the *D. melanogaster* JASPAR TFBS motifs (Sandelin *et al*, 2004) using FIMO (v5.5.5, default parameters) (Grant *et al*, 2011). The TFs were filtered for somatic, germline, or whole-ovary expression based on processed, publicly available single-cell RNA-seq data (Madrigal *et al*, 2026) accessed on https://wiki.flybase.org/wiki/FlyBase:Downloads_Overview.

### Screen for transcription factors that regulate TE niche using logistic regression

We predicted the *ENV* status based on the presence/absence of TFBS motifs using a generalised linear mixed effect model (GLMM) with a binomial error distribution and a logit link function with the *glmer()* function from the lme4 (v1.1-37) (Bates *et al*, 2015) in R (v4.5.1). Presence of a TFBS motif was treated as a fixed effect and TE identify was included as a random intercept. The bound optimisation by quadratic approximation (bobyqa) optimiser was used with the iteration limit increased to 100,000. Raw *p*-values were adjusted to control for the false discovery rate using the Benjamini-Hochberg procedure.

### TFBS co-occurrence analysis

The co-occurrence of TFBS in LTR regions of TEs was computed in R (v4.5.1) using the Jaccard Index, defined as:

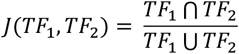

### Mapping *D. melanogaster* transcription factors to drosophilid reference genomes

To identify transcription factors of interest in the 249 fly genomes, the longest isoform from *Drosophila melanogaster* was obtained from FlyBase (https://flybase.org/) and mapped to the genomes using miniprot (v0.18-r281) (Li, 2023) with the -I flag to automatically select the maximum intron size based on genome length and -gtf. gffread (v0.12.7) (Pertea & Pertea, 2020) was then used to obtain the coding (-w) and protein (-y) sequences. Finally, search36 (v36.3.8g) (Pertea & Pertea, 2020) was used to obtain the SW score between the *D. melanogaster* protein sequences and the ones identified in other species. The highest SW score was kept for each species in cases where multiple hits were found. Similarity values <50% were discarded and the final SW scores were normalised to the length of the corresponding protein in *D. melanogaster*.

## Code availability

The scripts required to implement the reported transposon annotation pipeline are available on GitHub (https://github.com/annamaria23/retrotransposon-atlas) (Papameletiou & Bornelöv, 2026).

## Data availability

The curated transposon libraries and their metadata are available on GitHub (https://github.com/annamaria23/retrotransposon-atlas) (Papameletiou & Bornelöv, 2026).

## Acknowledgements

This work is dedicated to our colleague, mentor and supervisor, Gregory J. Hannon, who passed away in April 2026. We thank members of the Hannon and Bornelöv labs for fruitful discussions, and Peter Ziribagwa Sabakaki for initial help with phylogenetic analyses. We thank Anna Protasio for advice on using HMMs for TE detection. We thank the Scientific Computing core at the CRUK Cambridge Institute for HPC resources.

## AUTHOR CONTRIBUTIONS

All authors conceived the study. A-M.P. performed all investigations. A-M.P. annotated TEs with assistance from S.B.. A-M.P., S.B., B.C.N., and G.J.H. interpreted the results. S.B., B.C.N., G.J.H. supervised the work. A-M.P., S.B., B.C.N. and G.J.H. wrote the manuscript.

