## Supplementary Figure S1-S20 for "Lifestyles of *Gypsy*-family transposons shape their regulatory mechanisms"

Affiliations:
[1]Cancer Research UK Cambridge Institute, University of Cambridge, Cambridge, UK
[2]Department of Biochemistry, University of Cambridge, Cambridge, UK

Keywords:
Transposable elements, *Drosophila*, transcriptional regulation

**Supplementary Figures**

**Figure S1:** Quality control and genome selection
**Figure S2:** Transposable element content across the 249 selected drosophilid genomes
**Figure S3:** TE profiles of 249 drosophilid species
**Figure S4:** Representation of the TE annotation pipeline
**Figure S5:** Pipeline benchmarking
**Figure S6:** LTR identity for annotated TE consensus sequences
**Figure S7:** Small RNA sequences indicate that TEs are potentially under piRNA control
**Figure S8:** *Gypsy* TEs are likely targeted by the piRNA pathway
**Figure S9:** Presence of Gypsy-family LTR retrotransposons
**Figure S10:** Unrooted *POL* phylogenetic trees of all TEs across 249 drosophilid species
**Figure S11:** Divergence metrics for TEs across species
**Figure S12:** Full length insertions of TEs across species
**Figure S13:** TE genetic divergence landscapes across the species
**Figure S14:** Full-length TE genetic divergence landscapes across the species
**Figure S15:** Location preferences for TEs across species
**Figure S16:** Extensive phylogenetic incongruencies across TE evolutionary histories
**Figure S17:** HTT of TEs across drosophilid genomes
**Figure S18:** Number of motifs in each LTR
**Figure S19:** Distribution of motif scores for the top-scoring motif in each LTR
**Figure S20:** Conservation of six transcription factors predicted to be associated with germline or somatic TE expression across species

**Supplementary Tables**

**Table S1** (separate file). Table summarising drosophilid species names and percentages of each TE superfamily, as outputted by EDTA, RepeatMasker and RepeatModeler.
**Table S2** (separate file). Table summarising the drosophilid sRNA-seq libraries used in this study.
**Table S3** (separate file). Table summarising the drosophilid genome assemblies used in this study.

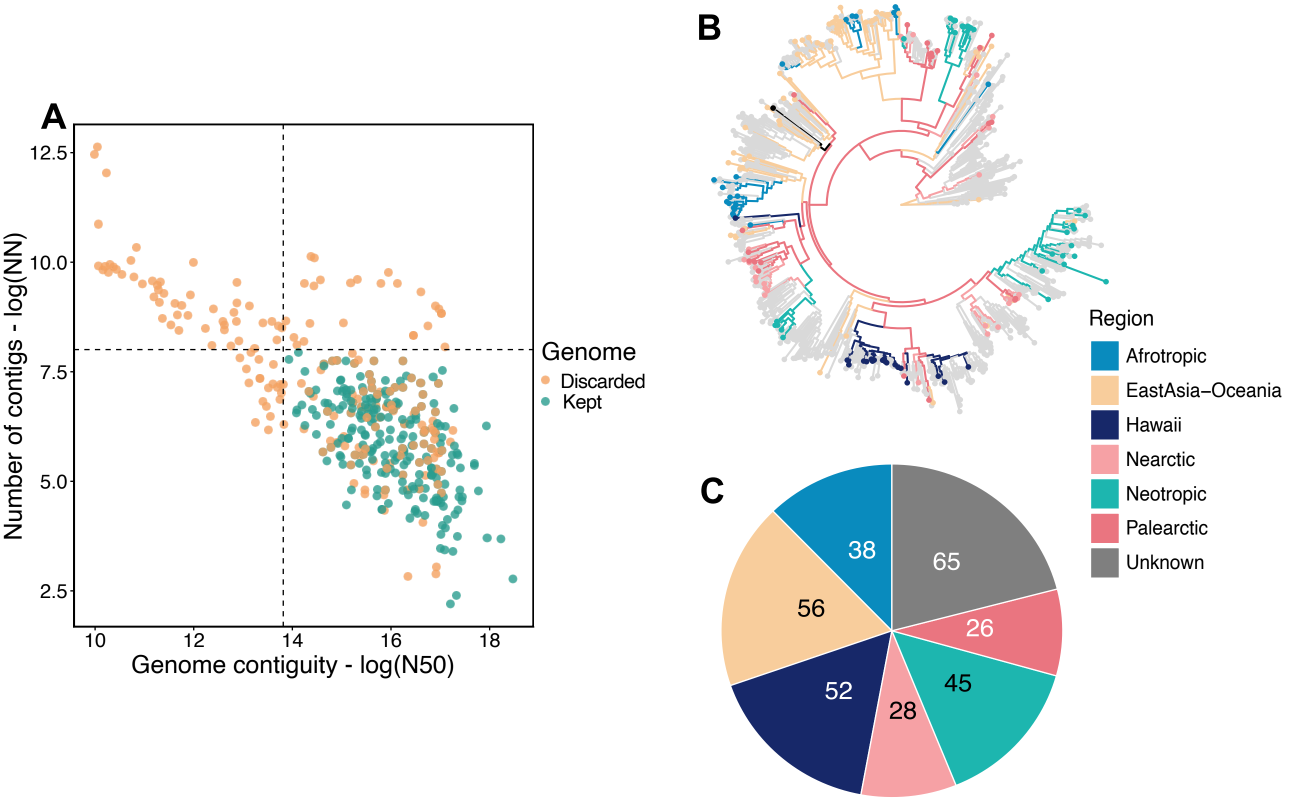


**Figure S1. Quality control and genome selection**

**A:** A scatterplot of quality control metrics (NN and N50), with the species which passed the control filters highlighted (green). Where there were multiple genome assemblies for the same species, the highest-quality assembly was selected (see Methods for more details). **B:** Displayed is a phylogenetic tree of 450 drosophilid species with the species used in this study highlighted and coloured by geographic region. **C**: The pie chart shows the geographical distributions of species harbouring the genomes used in this study. The geographic regions of drosophilid species were obtained from (Pianezza & Kofler, 2025).

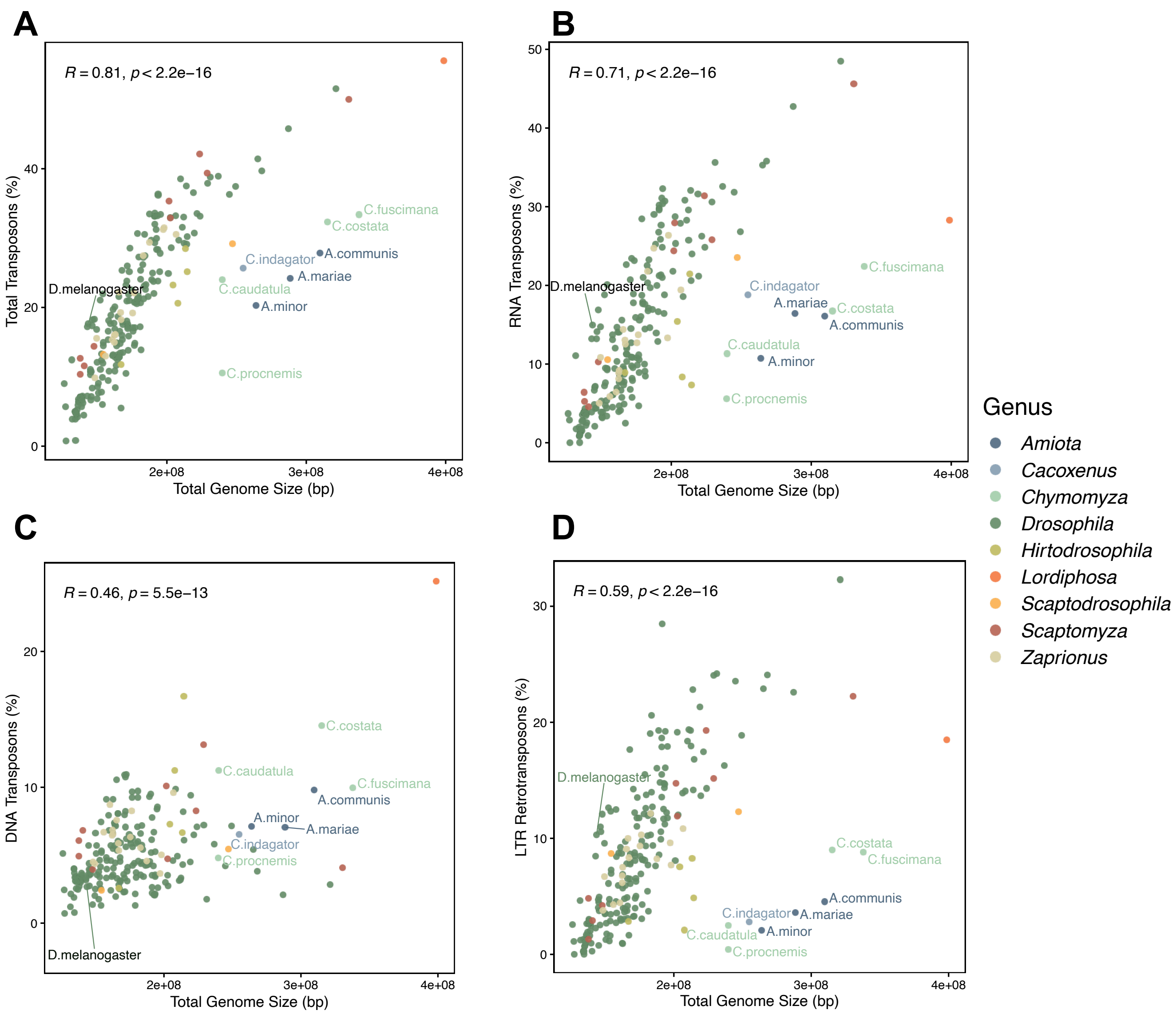


**Figure S2. Transposable element content across the 248 selected drosophilid genomes**

**A-D:** Scatterplots show total genome size against the total nucleotide coverage of all transposable elements (A), RNA (B) and DNA (C) and LTR retrotransposons (D).

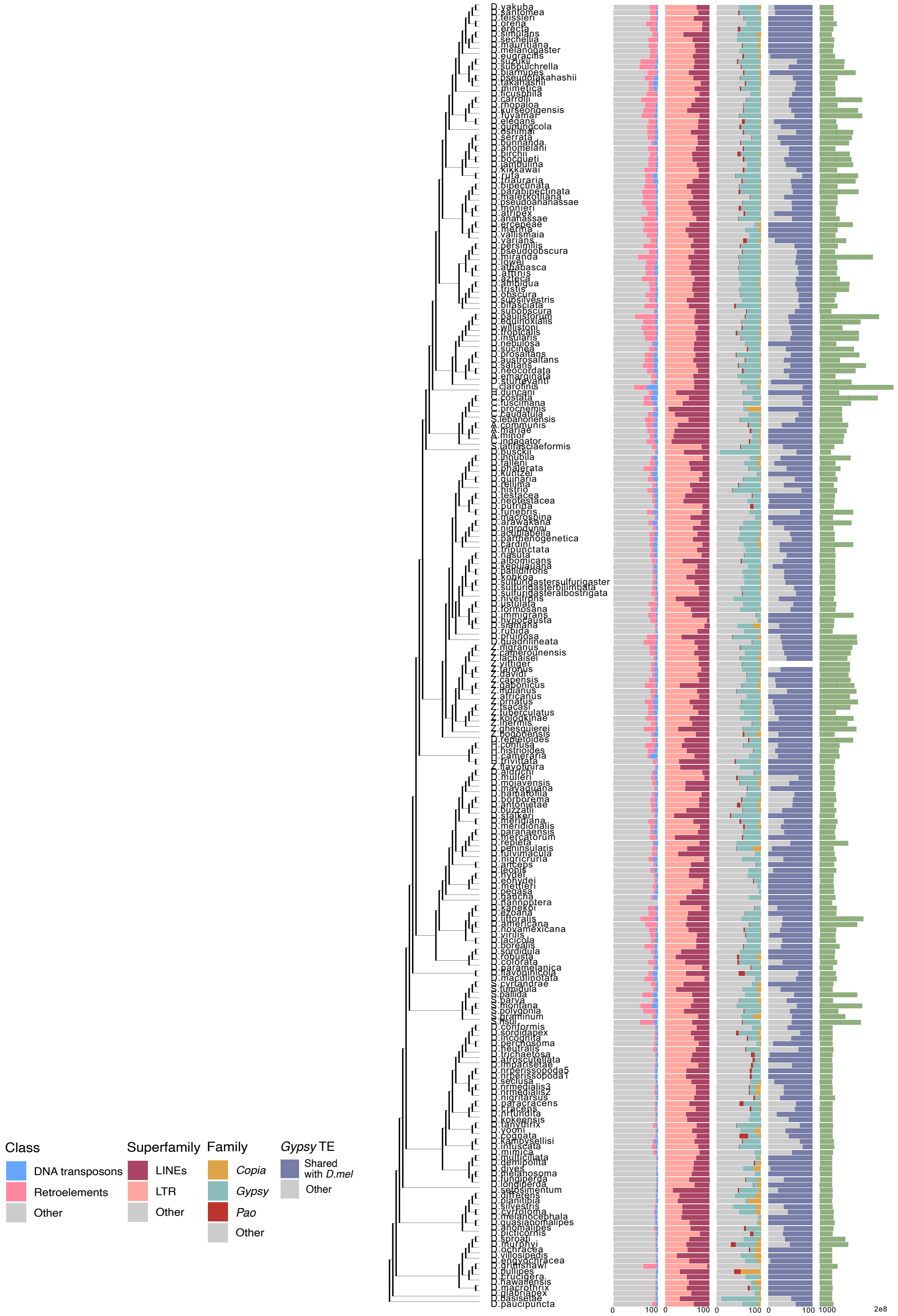


**Figure S3: TE profiles of 248 drosophilid species**

Shown is a phylogenetic tree of 248 drosophilid species, the percentages of TE class, Retrotransposon Superfamily, and LTR Family for each species as well as the percentage of *D. melanogaster*-like *Gypsy*-family TEs, obtained by the EDTA and RepeatMasker tools.

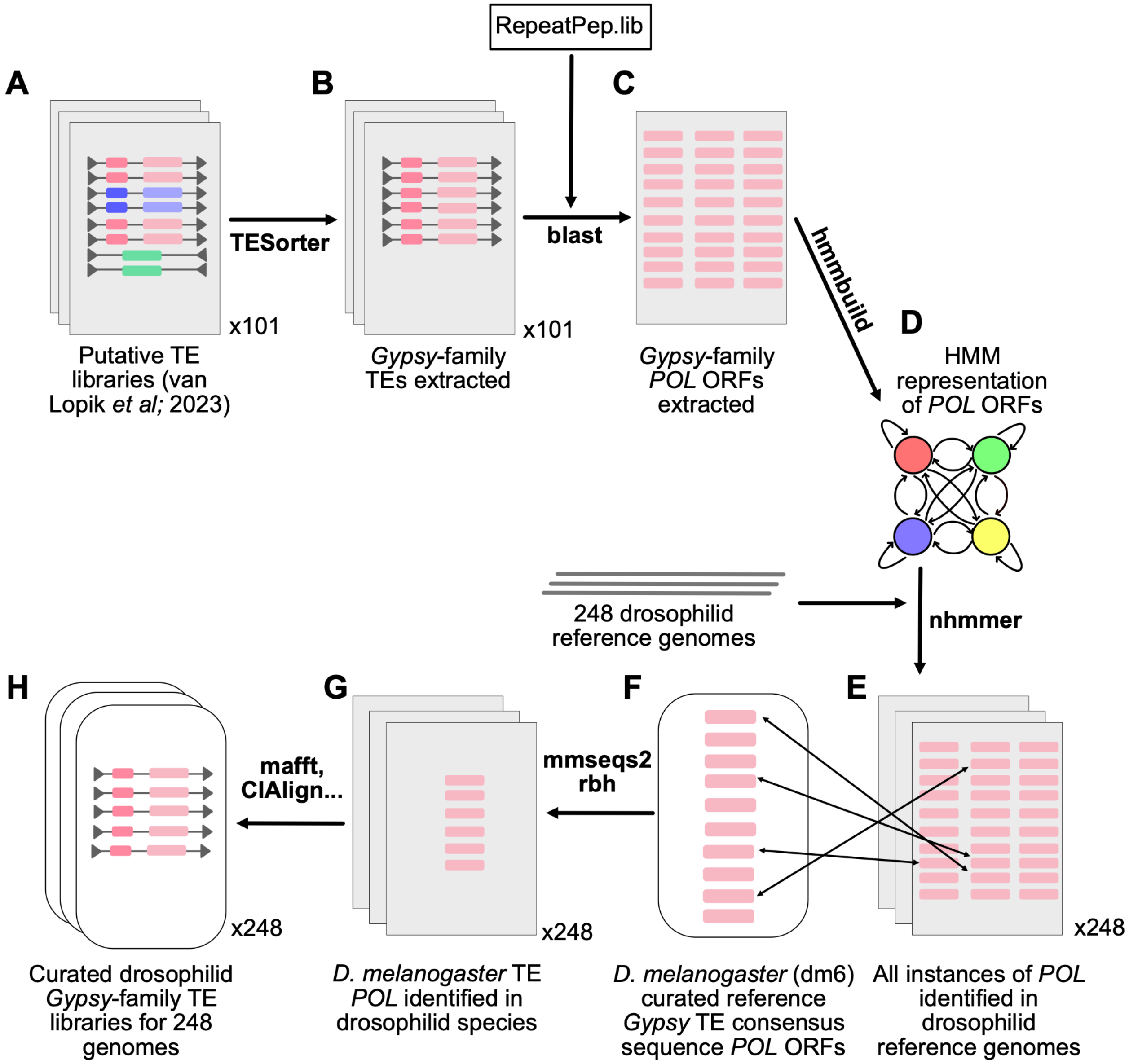


**Figure S4: Representation of the TE annotation pipeline**

**A**: Previously generated TE libraries (van Lopik *et al*, 2023) from 101 drosophilid genomes were used as a starting point. **B:** *Gypsy*-family LTR TEs were extracted from the TE libraries using TESorter. **C**: The TE *POL* open reading frames were extracted by using *blast* against the *POL* peptide database from RepeatMasker (RepeatPep.lib). **D-E:** The *POL* ORFs were used to generate a HMM representation of the *POL* protein using hmmer (D) which was then used to identify all instances of the *POL* protein in each genome (E). **F:** *mmseqs2* reciprocal best hits were then used to assign the *POL* protein the best match from the *D. melanogaster* consensus *POL* proteins. **G:** These sequences were then converted to a consensus sequence by a *blast* search against the genome to get the top 15 matching sequences, extended and then turned into consensus sequences using *mafft* and *ClAlign*.

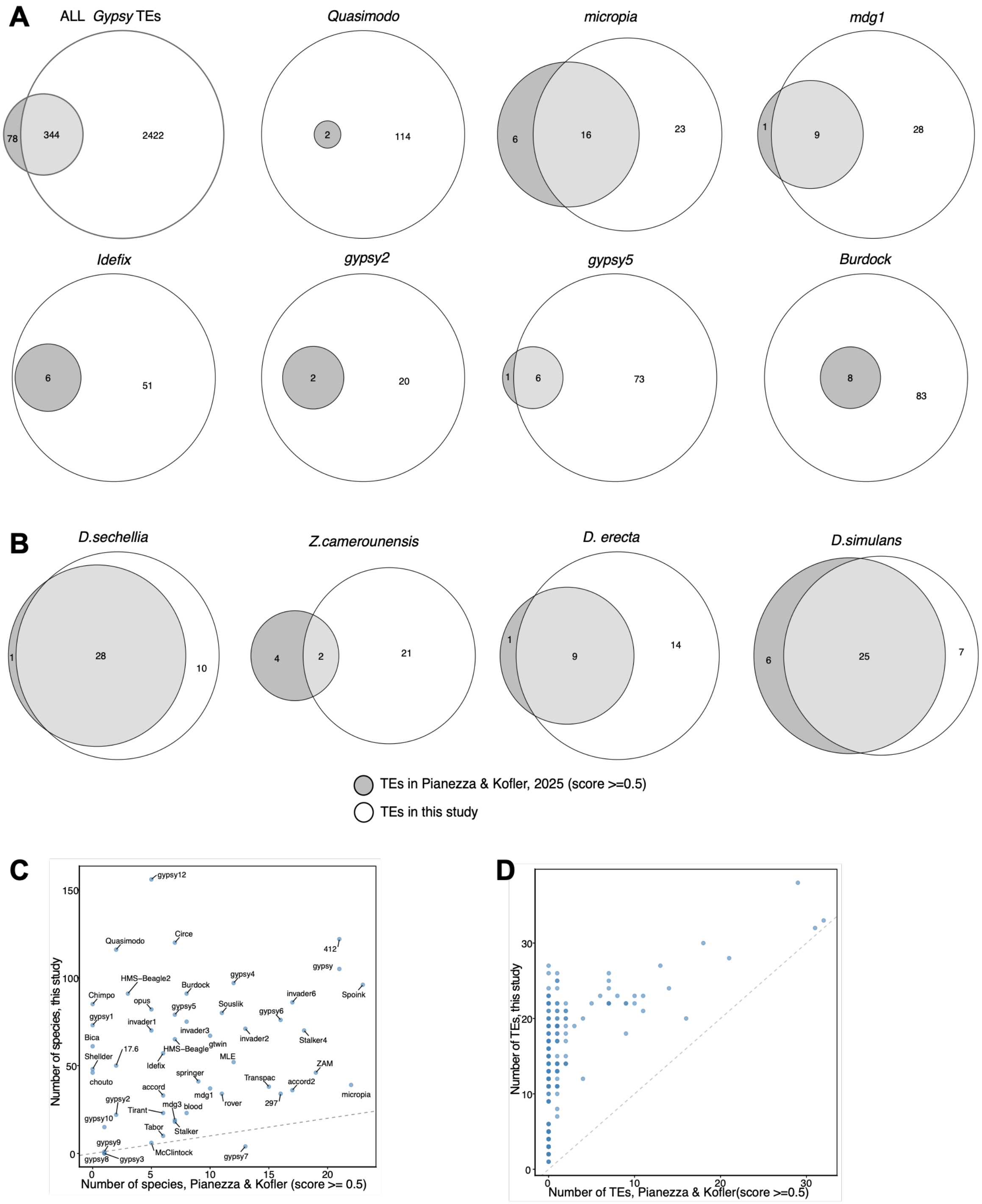


**Figure S5: Pipeline benchmarking**

**A:** Venn diagrams of species with the indicated TEs identified across our study and Pianezza and Kofler 2025. Shown are results for all *Gypsy*-family TEs combined (top left) and seven representative families. **B:** Venn diagrams of distinct *D. mel*-like *Gypsy*-family TEs identified across four representative species. **C**: Comparison of the number of species in which each *D. mel*-like *Gypsy* TE was annotated in Pianezza and Kofler 2025 versus our study. **D**: Comparison of the number of *D. mel*-like *Gypsy* TEs annotated in each species (excluding *D. melanogaster*) in Pianezza and Kofler 2025 versus our study.

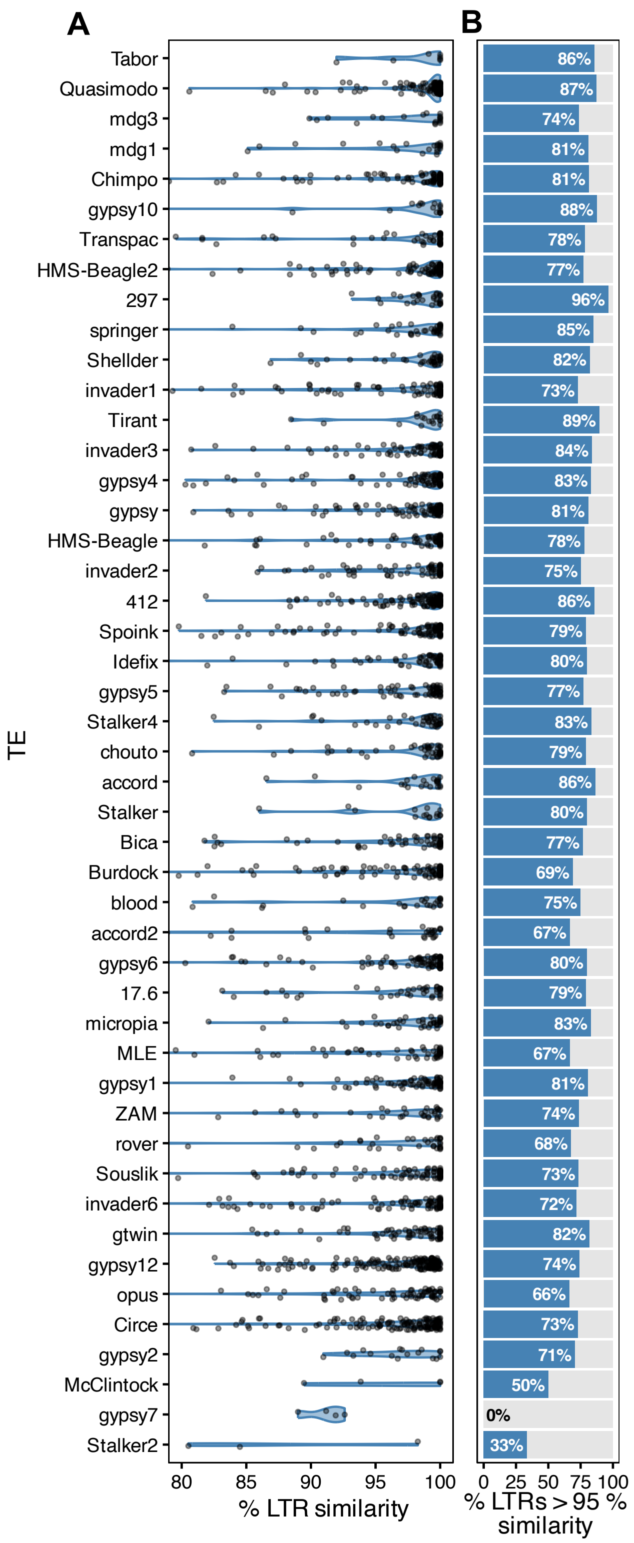


**Figure S6: LTR identity for annotated TE consensus sequences**

**A**: Violin plots show the distribution of identity (obtained through *blast*), of the 3' and 5' LTRs of TE consensus sequences. **B**: A bar plot shows the percentage of *Gypsy* TE consensus sequences for which the 5' and 3' LTRs share >= 95% identity, for each individual TE.

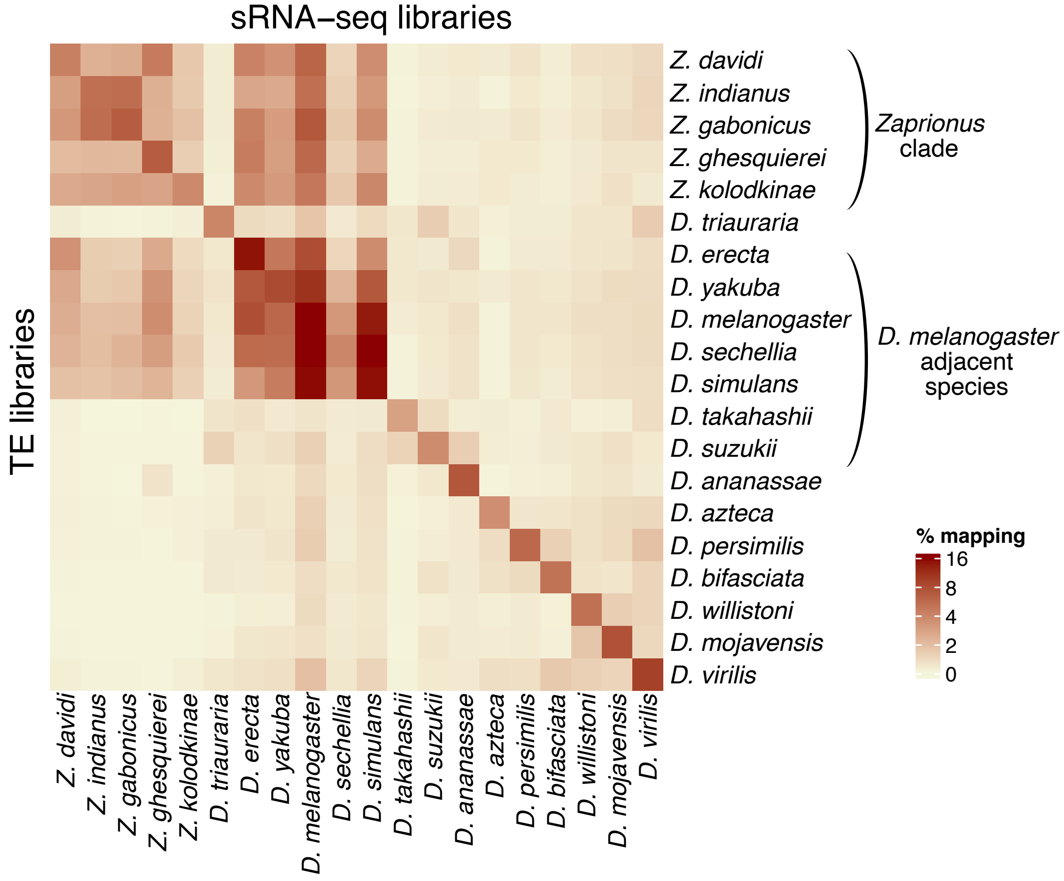


**Figure S7: Small RNA sequences indicate that TEs are potentially under piRNA control**

The heatmap shows the $\log_2$ (mapping percentage) of sRNA-seq libraries from 15 *Drosophila* species, arranged according to their order in the species phylogeny, to 22 Drosophilid species. In cases where multiple sRNA-seq libraries are available for a single species, an average of available libraries is shown.

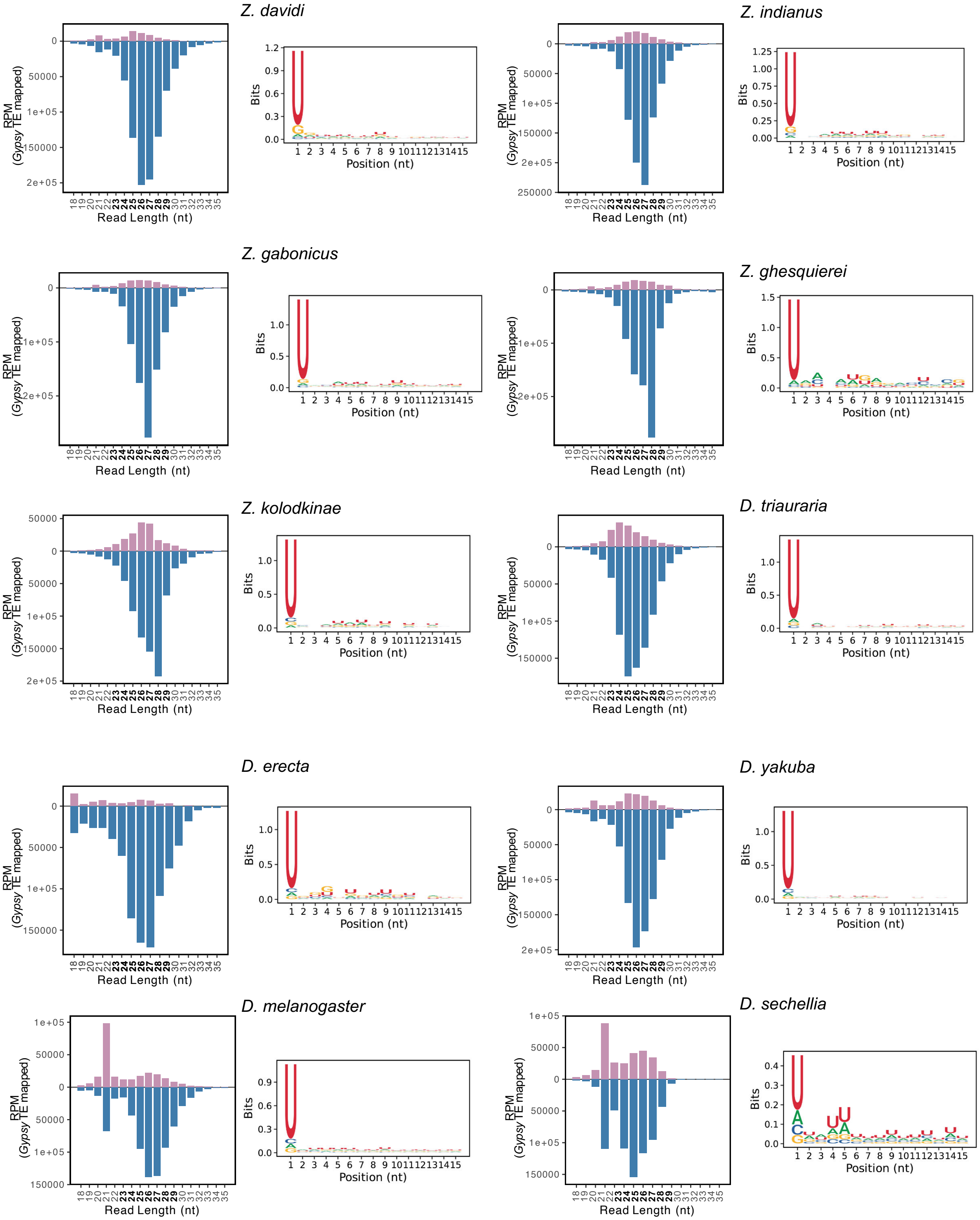
Z. davidi
Z. indianus
Z. gabonicus
Z. ghesquierei
Z. kolodkinae
D. triauraria
D. erecta
D. yakuba
D. melanogaster
D. sechellia
RPM
(Gypsy TE mapped)
Read Length (nt)
Bits
Position (nt)

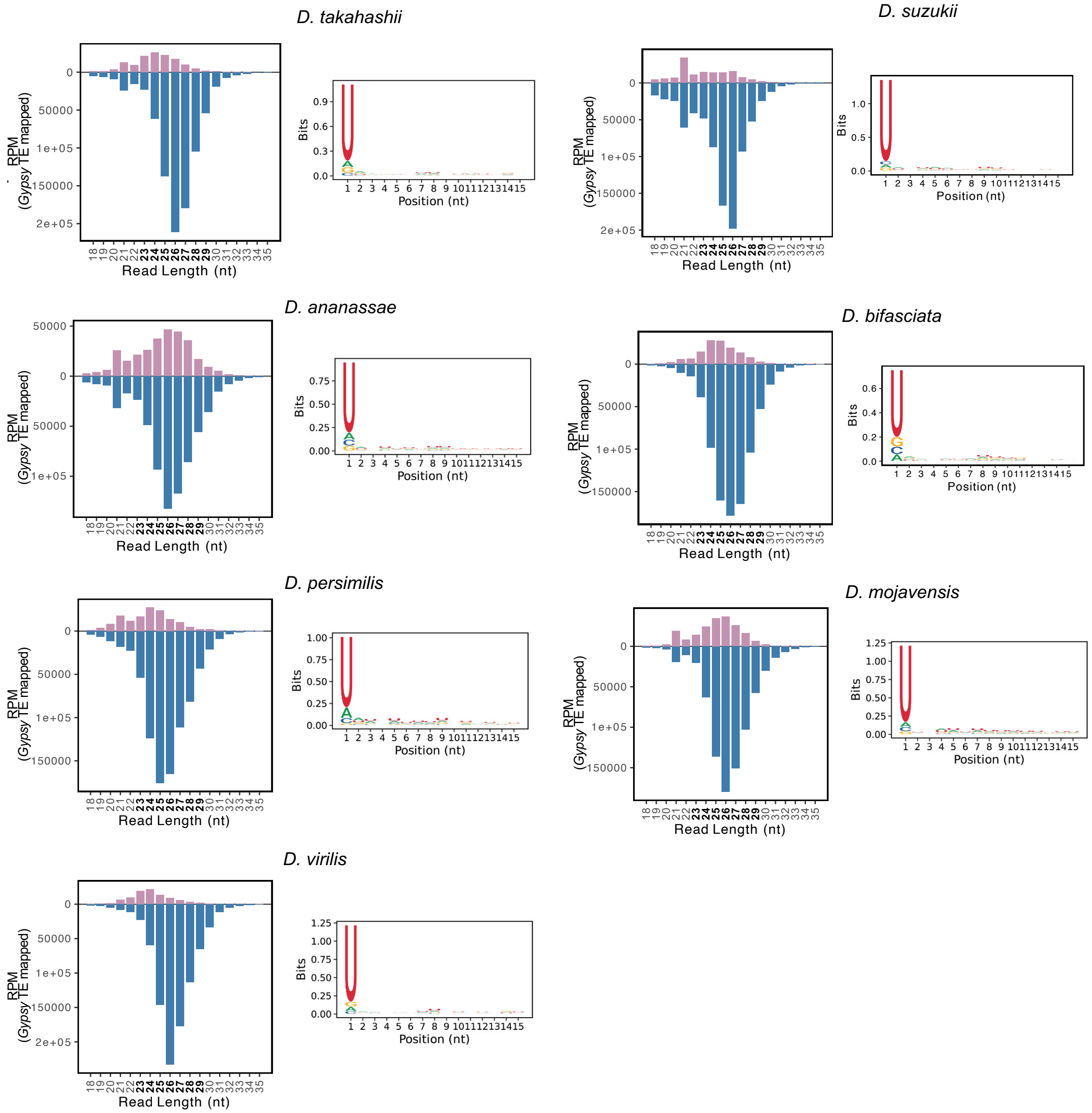


**Figure S8: *Gypsy* TEs are likely targeted by the piRNA pathway**

Bar plots show the length distribution of piRNAs mapping to the species' own TE libraries in the sense and antisense direction and sequence logo plots show the nucleotide frequency of 15 bp at the 5' ends of piRNAs for one representative sRNA-seq library for each species.

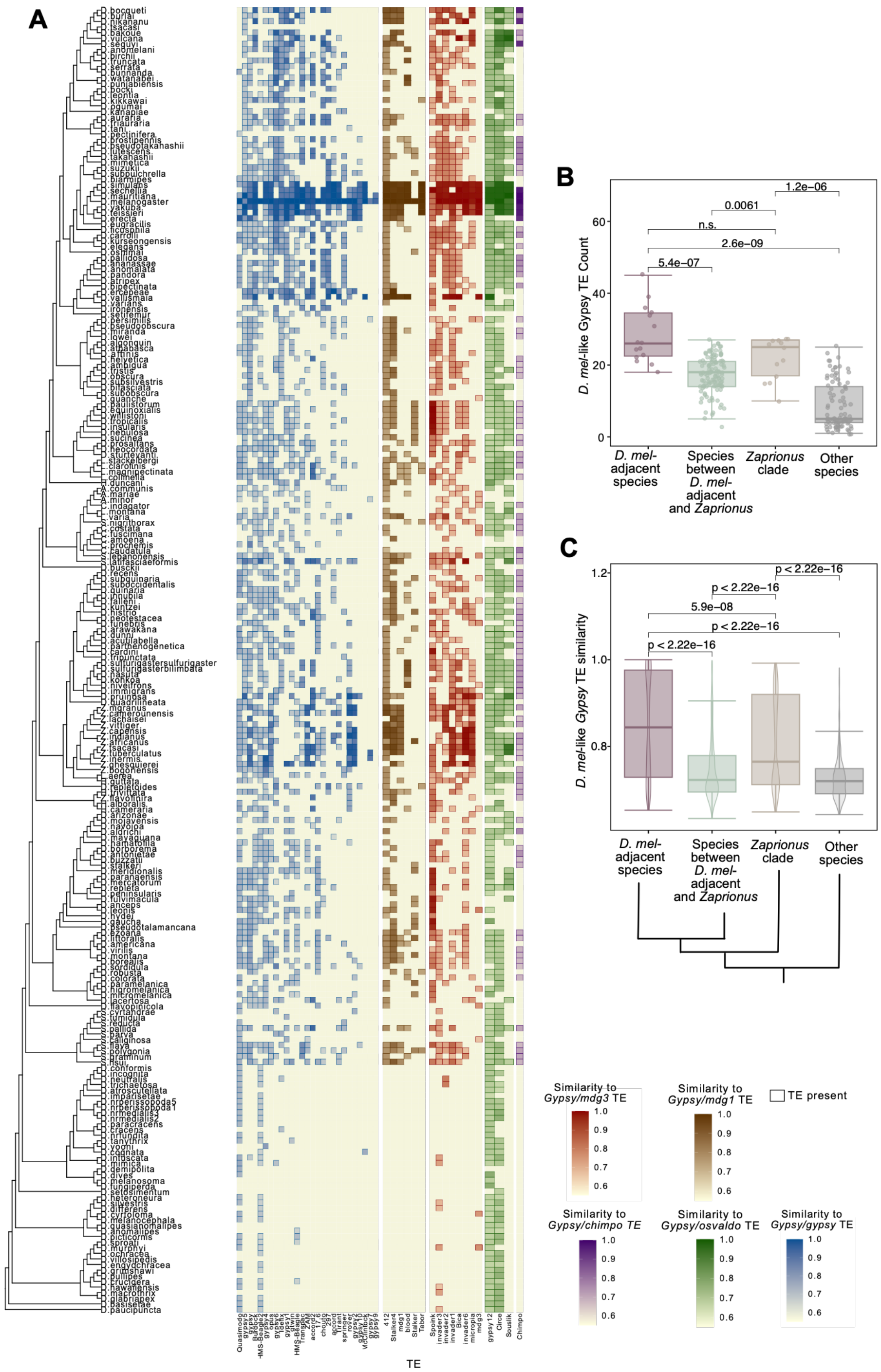


**Figure S9: Presence of Gypsy-family LTR retrotransposons**

**A:** The phylogenetic tree and heatmap show TE presence (≥70% similarity) and absence across 248 species. Heatmap colours indicate the similarity to the corresponding *POL* ORF in *D. melanogaster*. **B**: Boxplot showing the distribution of the number of *D. mel*-like TEs identified by our pipeline for species in the *D. melanogaster*-adjacent species(*D. yakuba* – *D. ficusphila*), *Zaprionus* genus, intermediate species bridging the two groups, and species located further away on the drosophilid phylogenetic tree. The p-values shown are a result of a Wilcoxon rank-sum test. **C**: Boxplots showing the distribution of the similarity of the of *D. mel*-like *Gypsy* TEs annotated in each species to the *D. melanogaster* reference POL sequences. The p-values shown are a result of a Wilcoxon rank-sum test. The cladogram shown is a schematic representation of the phylogenetic distances between the species in each group.

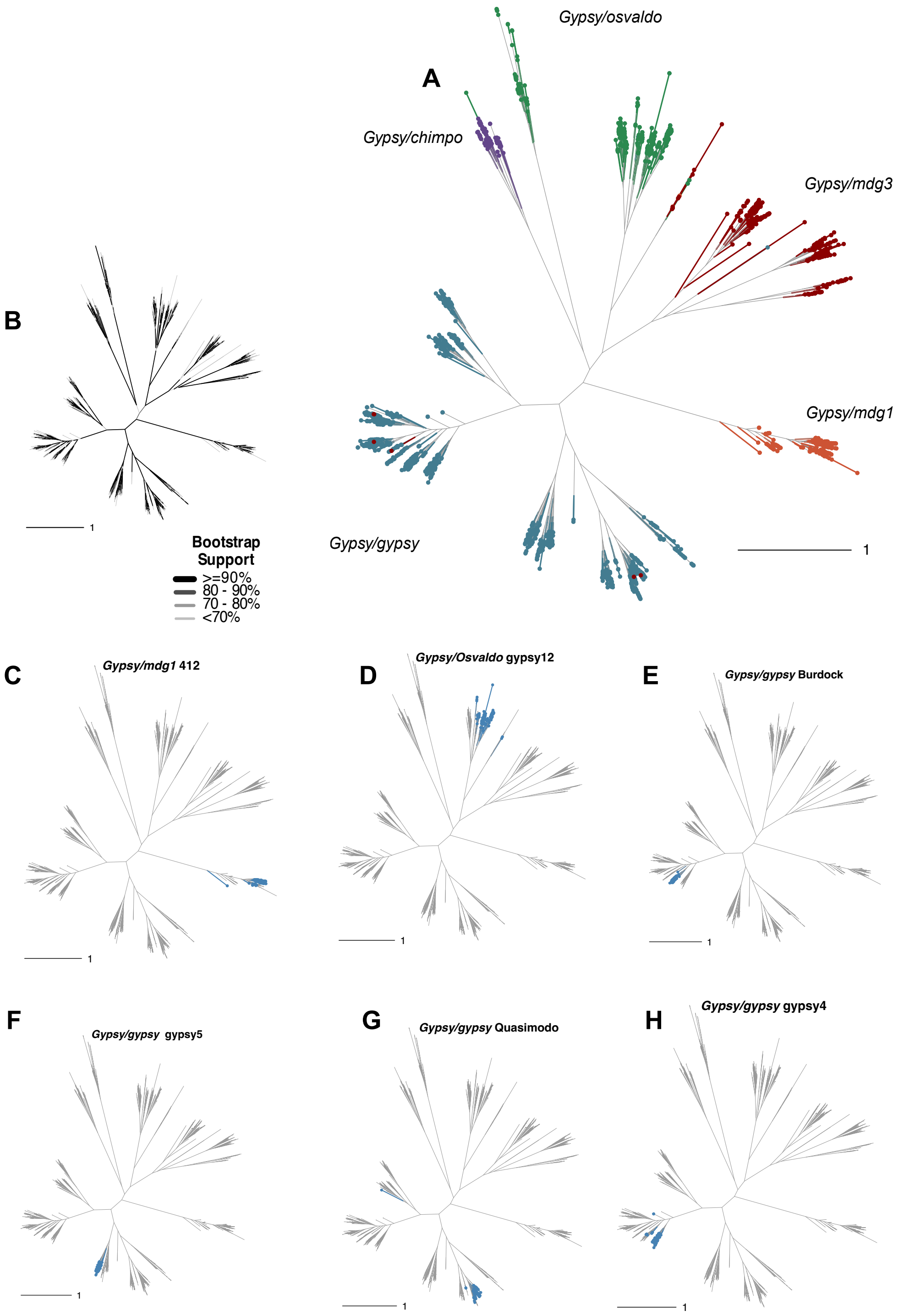


**Figure S10: Unrooted *POL* phylogenetic trees of all TEs across 248 drosophilid species**

**A**: Shown is an unrooted *POL* phylogenetic tree with the four main TE clades (*Gypsy/gypsy, Gypsy/mdg1, Gypsy/Osvaldo* and *Gypsy/mdg3*) coloured. The scale bar indicates amino acid substitutions per site. **B**: Unrooted *POL* phylogenetic tree with edge colours and widths determined by their bootstrap support values. **C-H**: Unrooted *POL* phylogenetic tree highlighting (blue) consensus sequences from different species for *412* (**C**), *gypsy12* (**D**), *Burdock* (**E**), *gypsy5* (**F**), *Quasimodo* (**G**), *gypsy4* (**H**) TEs. The TEs shown were selected because they were widely present in many species. The scale bars indicate amino acid substitutions per site.

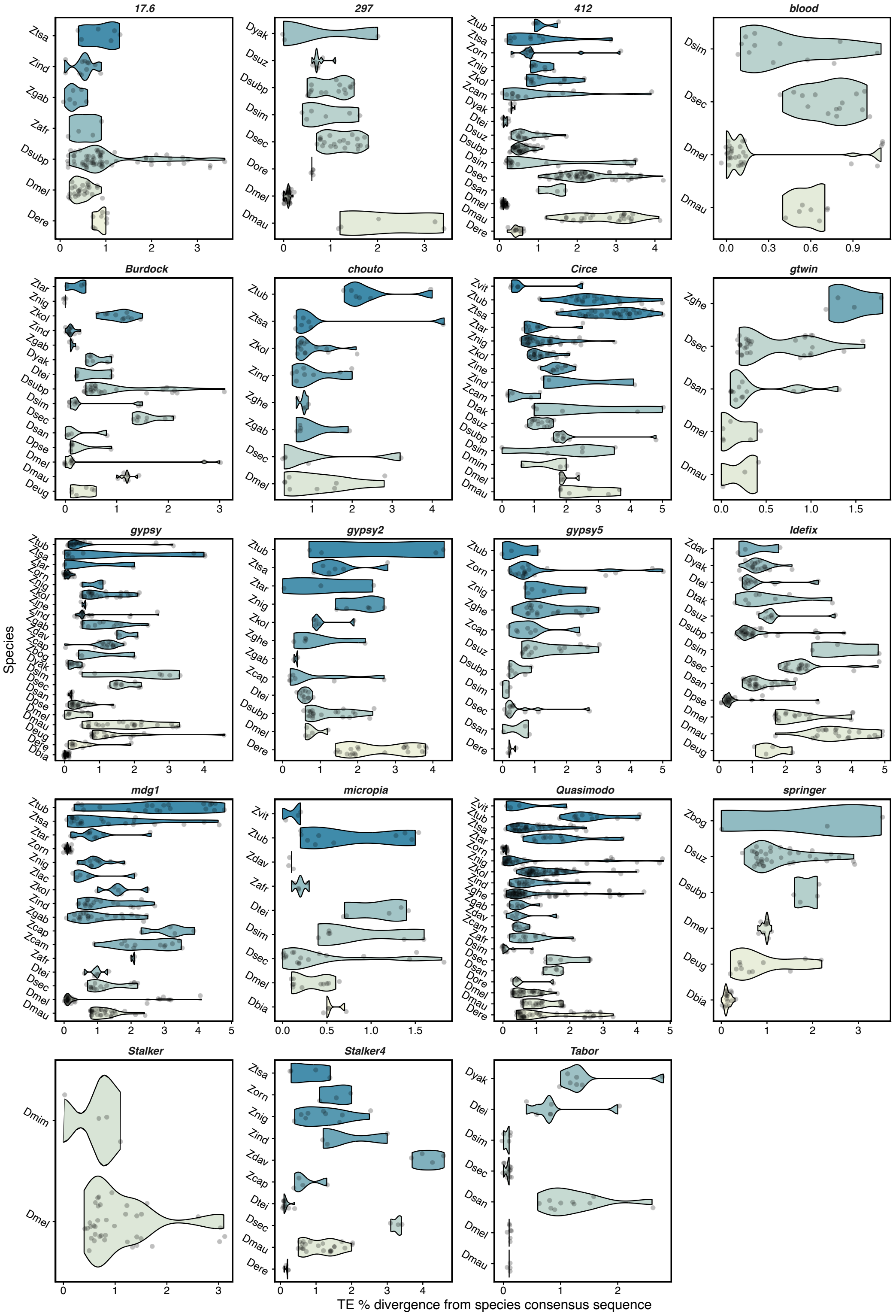


**Figure S11: Divergence metrics for TEs across species**

Ridge plots show distributions of the percent divergence from the consensus sequences for all full-length and high-quality genomic insertions of *Gypsy*-family LTR TEs (considered to be longer than 4,000 nucleotides and have a divergence to the consensus sequence lower than 5%) in the 16 melanogaster clade *Drosophila* species and 17 *Zaprionus* species used in this study.

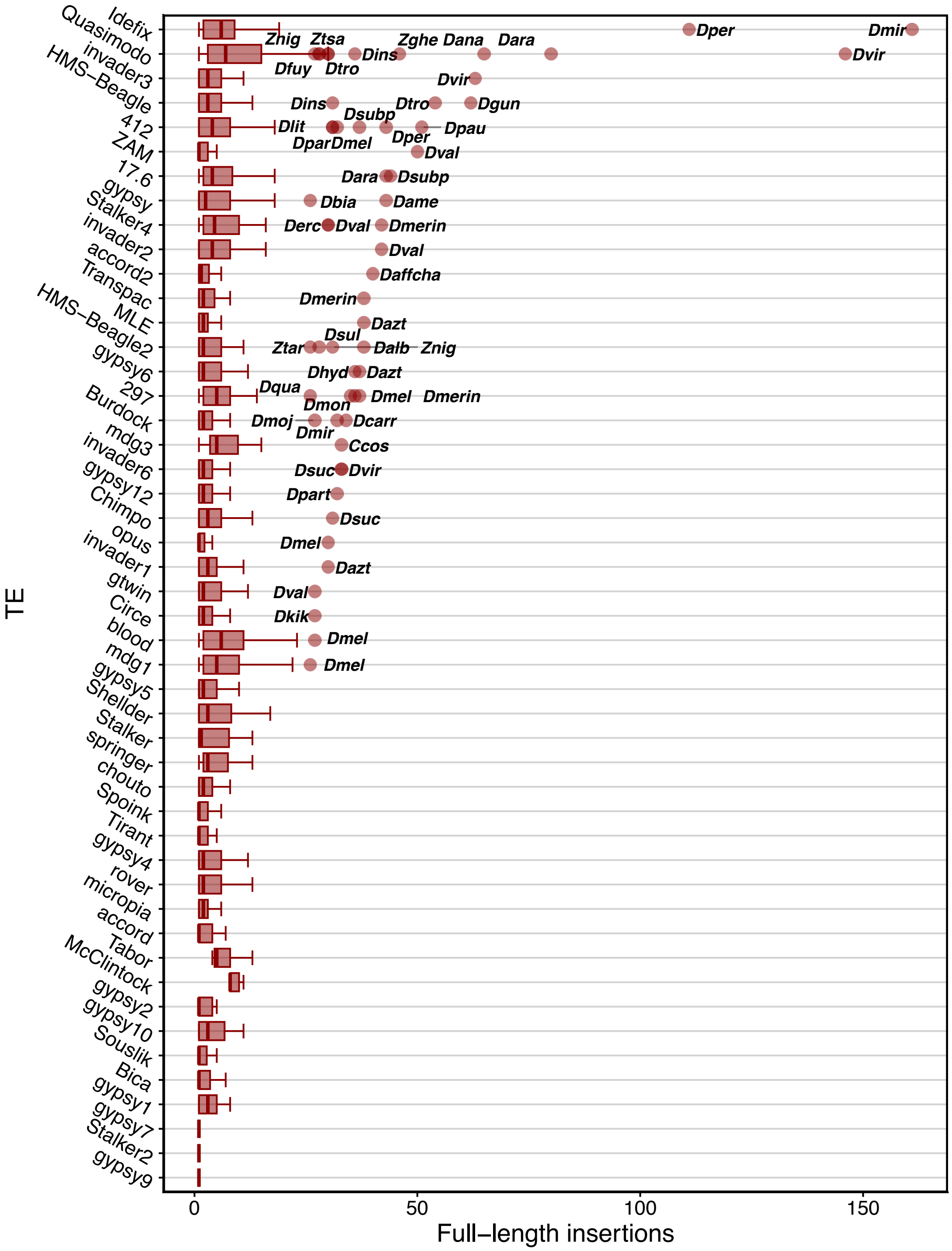


**Figure S12: Full length insertions of TEs across species**

Boxplots show the number of full-length and high-quality genomic insertions of *Gypsy*-family LTR TEs (considered to be longer than 4,000 nucleotides and have a divergence to the consensus sequence lower than 1%). Species with more than 25 detected insertions of a specific TE are highlighted.

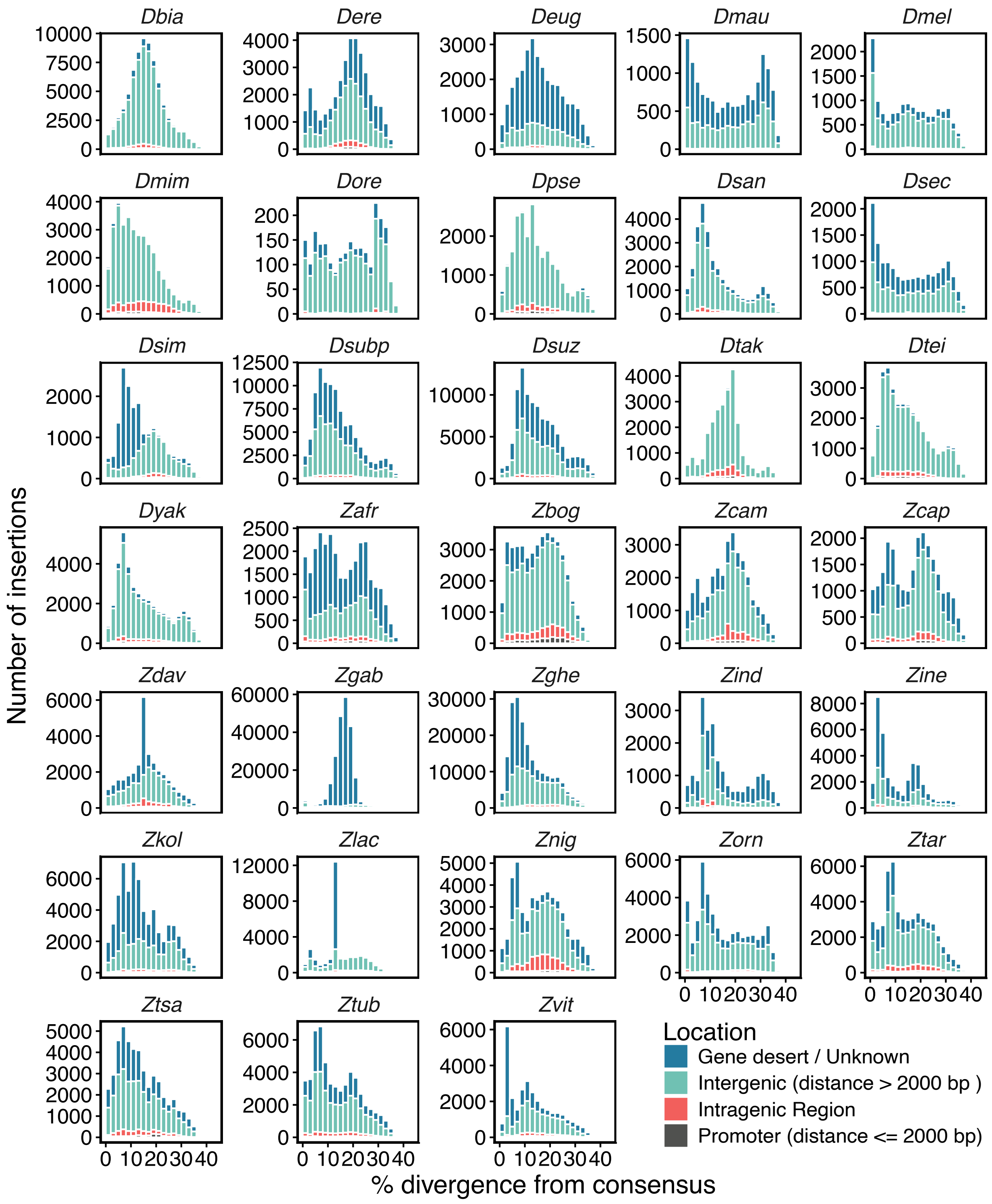


**Figure S13: TE genetic divergence landscapes across the species**

TE divergence landscapes of all genomic insertions of *Gypsy*-family LTR TEs in the 16 melanogaster clade *Drosophila* species and 17 *Zaprionus* species used in this study are shown. Distances from the nearest gene are defined as the distance to the nearest *Drosophilidae* BUSCO gene.

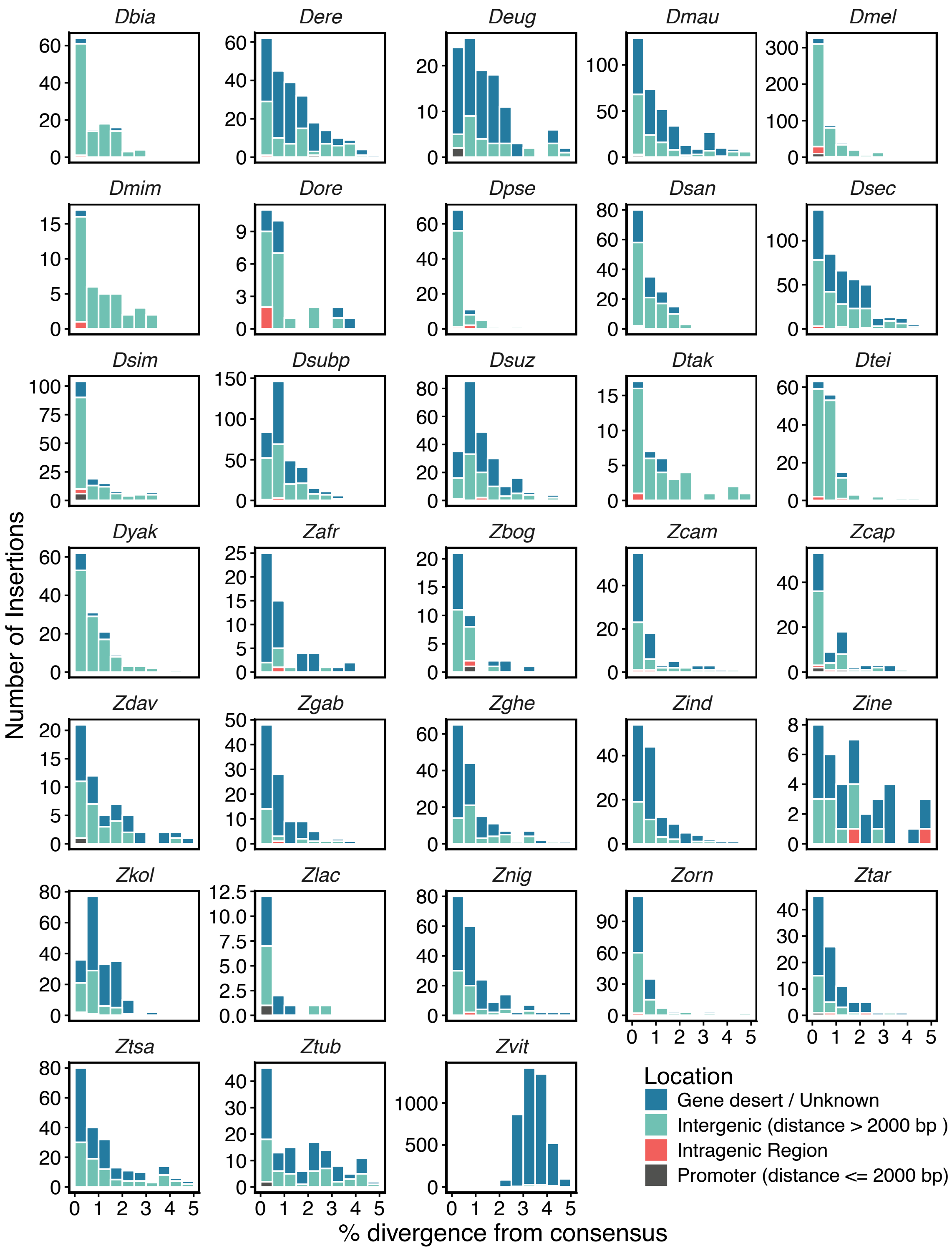


**Figure S14: Full-length TE genetic divergence landscapes across the species**

Shown are TE divergence landscapes of all full-length and high-quality genomic insertions of *Gypsy*-family LTR TEs (considered to be longer than 4,000 nucleotides and have a divergence to the consensus sequence lower than 5%) in the 16 melanogaster clade *Drosophila* species and 17 *Zaprionus* species used in this study. Distances from the nearest gene are defined as the distance to the nearest *Drosophilidae* BUSCO gene on the same strand as the TE insertion.

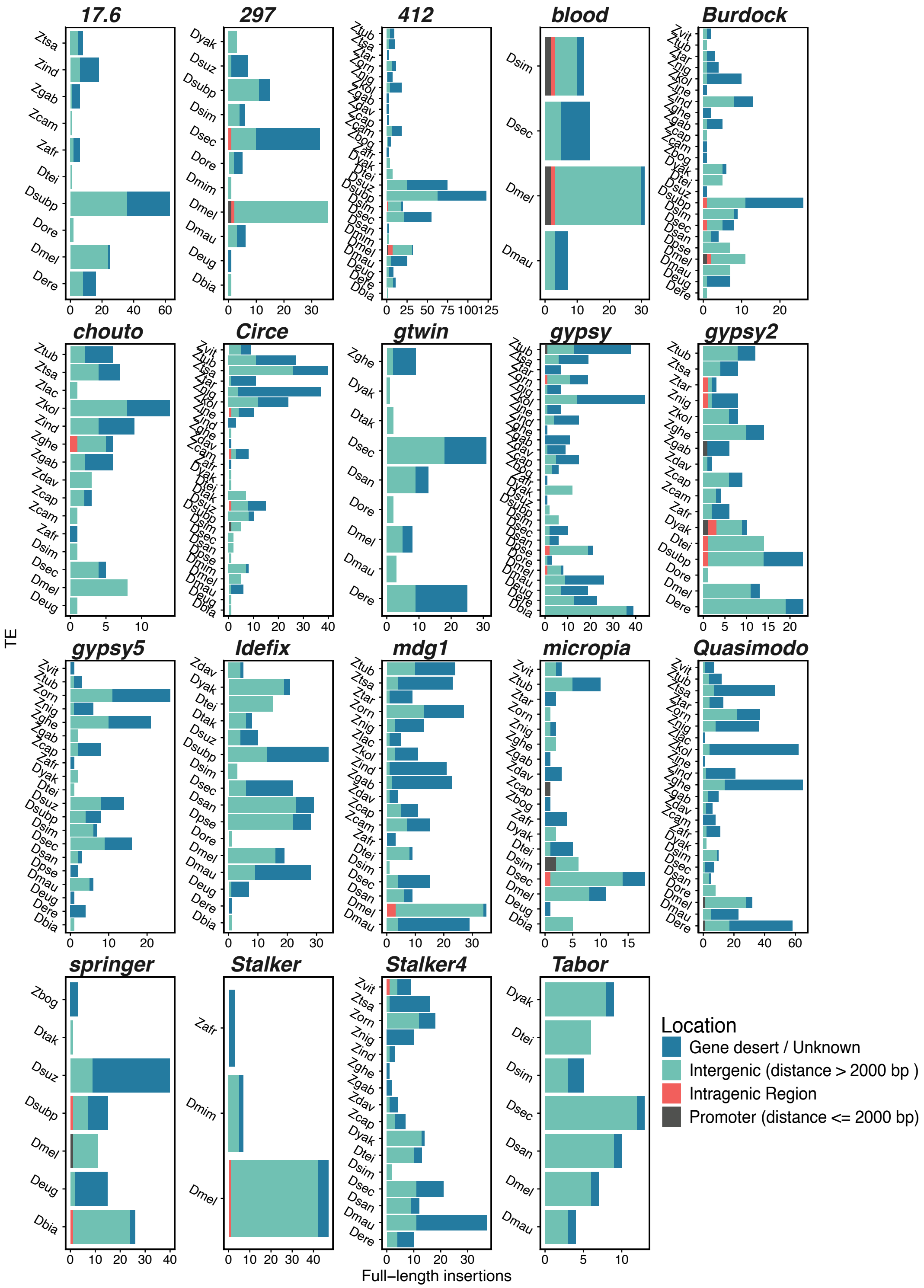


**Figure S15: Location preferences for TEs across species**

Bar plots show annotations of insertion locations of all full-length and high-quality genomic insertions of *Gypsy*-family LTR TEs (considered to be longer than 4,000 nucleotides and have a divergence to the consensus sequence lower than 5%) in the 16 melanogaster clade *Drosophila* species and 17 *Zaprionus* species used in this study. Distances from the nearest gene are defined as the distance to the nearest *Drosophilidae* BUSCO gene on the same strand as the TE insertion.

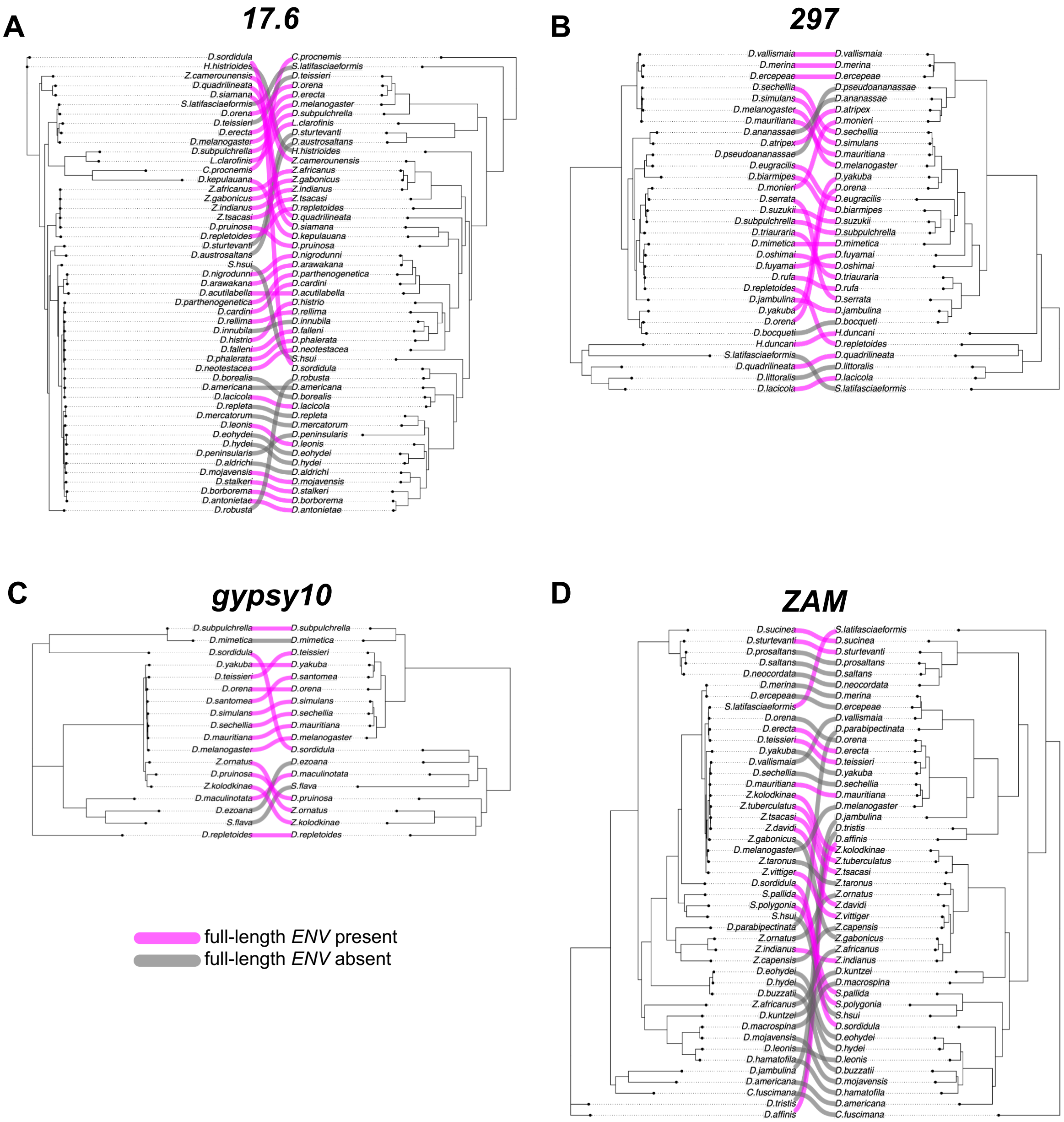


**Figure S16: Extensive HTTs across TEs' evolutionary history**

**A-E**: Displayed are phylogenetic tree tanglegrams showing the incongruences between the *POL* ORF phylogenetic tree (left) and the species phylogenetic tree (right) for *17.6* (A), *297* (B), *gypsy10* (C), and *ZAM* (D). Species in which the TE contains a full-length ENV ORF are coloured in pink.

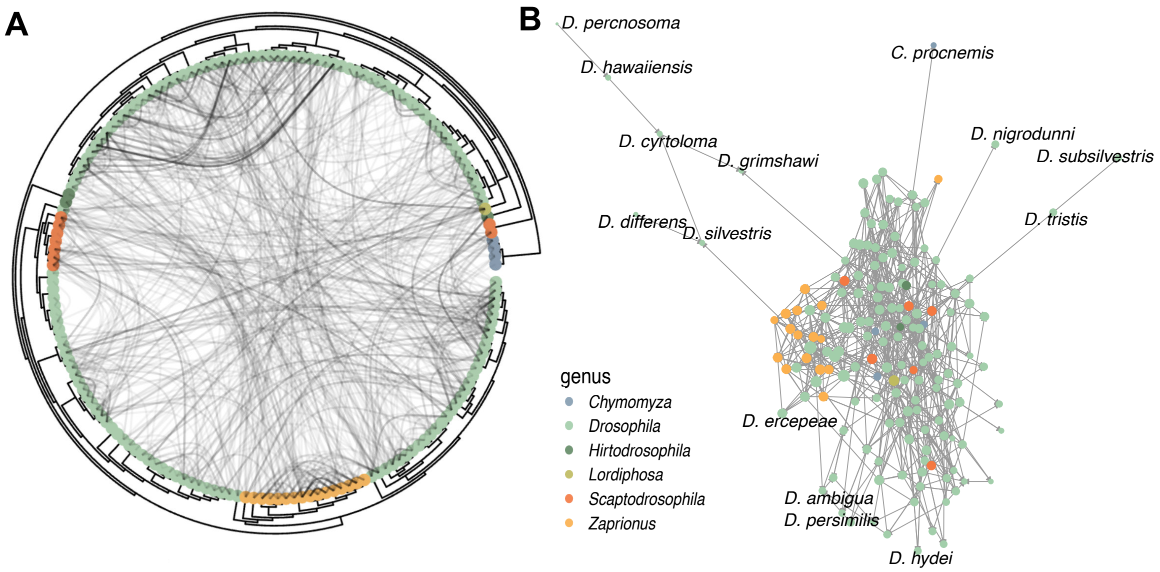


**Figure S17: Putative horizontal transfer of TEs across drosophilid genomes**

**A:** A phylogenetic tree is displayed with links between species where horizontal transposon transfer took place. **B**: HTT events between species are represented in a graph topology**.**

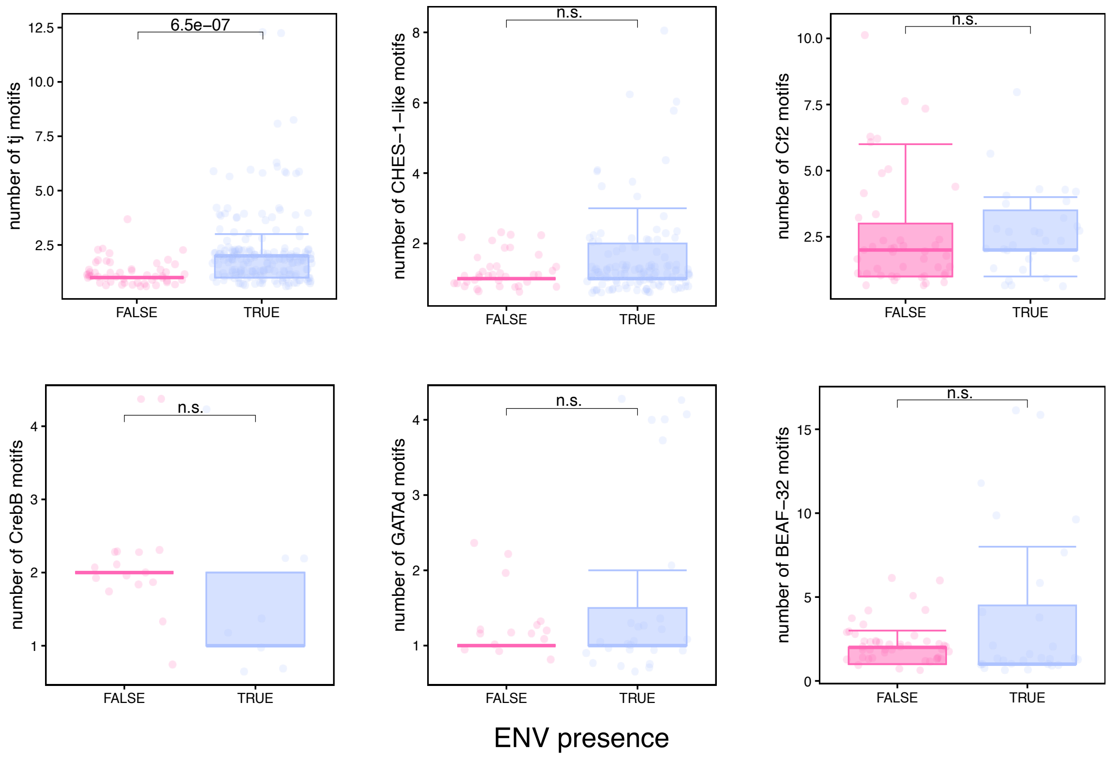


**Figure S18: Number of motifs in each LTR**

Displayed are boxplots showing the number of TF bidning motifs in the LTR of transcription factors significantly associated with TE expression in the germline (*CrebB, GATAd, Cf2, BAF-32*) and the soma (*CHES-1-like, tj*). The p-values are the results of a Wilcoxon rank-sum test.

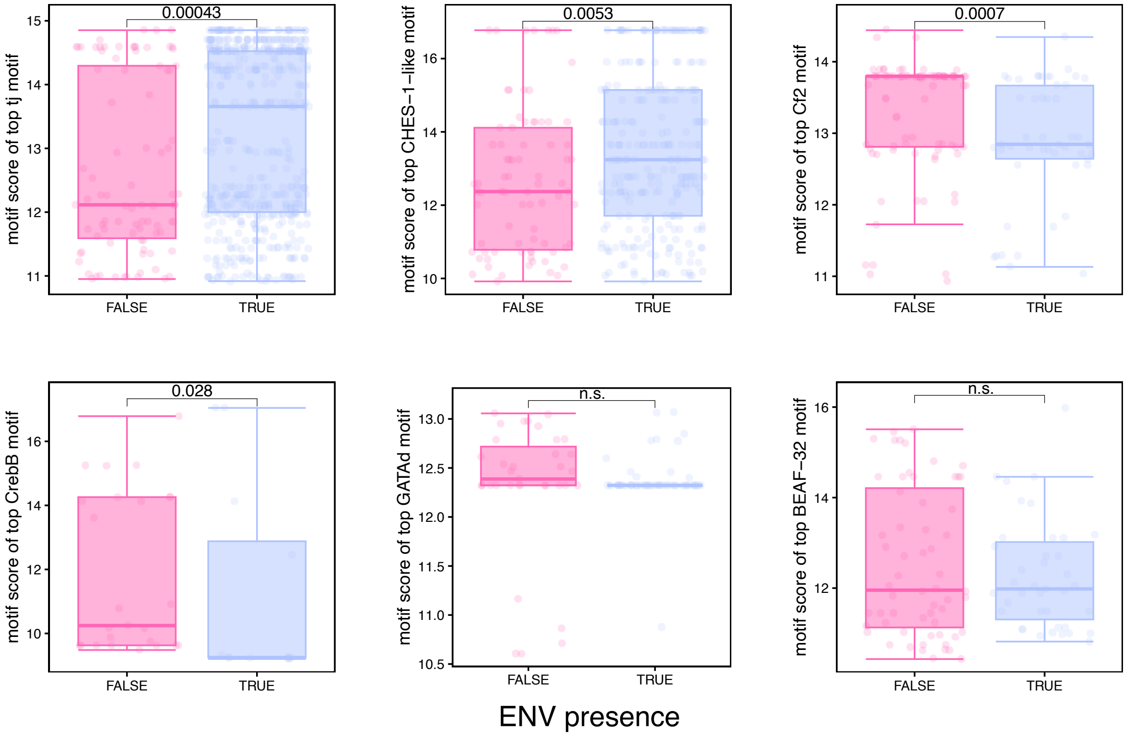


**Figure S19: Distribution of motif scores for the top-scoring motif in each LTR**

Displayed are boxplots showing the motif score (a.u.) for the top-ranking motif per LTR, outputted by the FIMO tool for TF bidning motifs in the LTR of transcription factors significantly associated with TE expression in the germline (*CrebB, GATAd, Cf2, BAF-32*) and the soma (*CHES-1-like, tj*). The p-values are the results of a Wilcoxon rank-sum test.

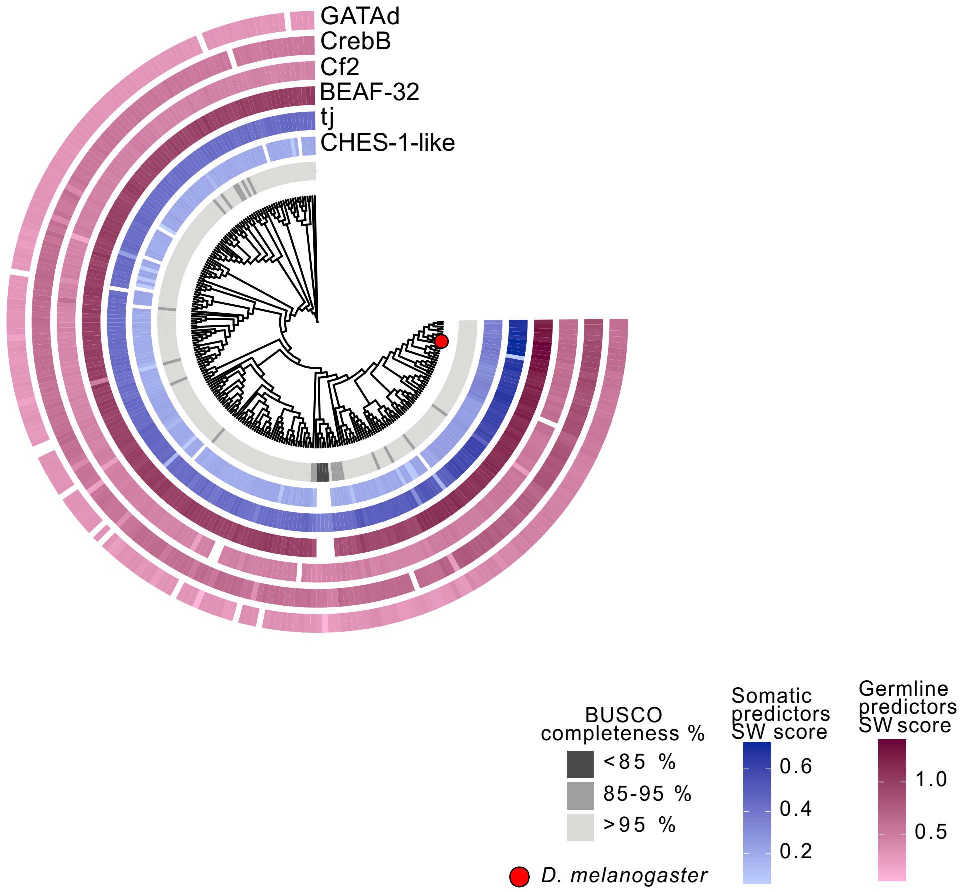


**Figure S20: Presence of six transcription factors predicted to be associated with germline or somatic TE expression across species**

The phylogenetic tree and heatmap show transcription factor presence and absence across 248 species for the transcription factors significantly associated with TE expression in the germline (*CrebB, GATAd, Cf2, BAF-32*) and the soma (*CHES-1-like, tj*). Heatmap colours indicate the Smith-Waterman alignment similarity to the protein sequence in *D. melanogaster*, normalised by the query (*D. melanogaster*) sequence.
